# RTL8 promotes nuclear localization of UBQLN2 to subnuclear compartments associated with protein quality control

**DOI:** 10.1101/2021.04.21.440788

**Authors:** Harihar Milaganur Mohan, Amit Pithadia, Hanna Trzeciakiewicz, Emily V. Crowley, Regina Pacitto, Nathaniel Safren, Chengxin Zhang, Xiaogen Zhou, Yang Zhang, Venkatesha Basrur, Henry L. Paulson, Lisa M. Sharkey

**Affiliations:** Department of Neurology, University of Michigan, Ann Arbor, MI 48109-2200; Graduate Program in Cellular and Molecular Biology, University of Michigan, Ann Arbor, MI 48109-2200; Department of Neurology, Northwestern University Feinberg School of Medicine, Chicago, IL 60611; Department of Computational Medicine and Bioinformatics, University of Michigan, Ann Arbor, MI 48109-2200; Department of Pathology, University of Michigan Medical School, Ann Arbor, MI 48109

**Keywords:** Ubiquilin, UBQLN2, RTL8, Nuclear Protein Quality Control, Ubiquitin Proteasome System

## Abstract

The brain expressed ubiquilins (UBQLNs) 1, 2 and 4 are a family of ubiquitin adaptor proteins that participate broadly in protein quality control (PQC) pathways, including the ubiquitin proteasome system (UPS). One family member, UBQLN2, has been implicated in numerous neurodegenerative diseases including ALS/FTD. UBQLN2 typically resides in the cytoplasm but in disease can translocate to the nucleus, as in Huntington’s disease where it promotes the clearance of mutant Huntingtin protein. How UBQLN2 translocates to the nucleus and clears aberrant nuclear proteins, however, is not well understood. In a mass spectrometry screen to discover UBQLN2 interactors, we identified a family of small (13 kDa), highly homologous uncharacterized proteins, RTL8, and confirmed the interaction between UBQLN2 and RTL8 both *in vitro* using recombinant proteins and *in vivo* using mouse brain tissue. Under endogenous and overexpressed conditions, RTL8 localizes to nucleoli. When co-expressed with UBQLN2, RTL8 promotes nuclear translocation of UBQLN2. UBQLN2 and RTL8 colocalize within ubiquitin-enriched subnuclear structures containing PQC components. The robust effect of RTL8 on the nuclear translocation and subnuclear localization of UBQLN2 does not extend to the other brain-expressed ubiquilins, UBQLN1 and UBQLN4. Moreover, compared to UBQLN1 and UBQLN4, UBQLN2 preferentially stabilizes RTL8 levels in human cell lines and in mouse brain, supporting functional heterogeneity among UBQLNs. As a novel UBQLN2 interactor that recruits UBQLN2 to specific nuclear compartments, RTL8 may regulate UBQLN2 function in nuclear protein quality control.

## Introduction

UBQLN2 is a member of the ubiquilin family of proteins that participate in ubiquitin-dependent protein quality control (PQC) pathways [1-3]. While UBQLN2 is best known as a shuttle adaptor protein for the ubiquitin-proteasome system (UPS), it is also implicated in autophagy, stress granule dynamics and chaperone-dependent pathways [4-7]. This versatility reflects various functional domains in UBQLN2, including its ubiquitin-binding (UBA) and ubiquitin-like (UBL) domains, as well as its propensity to engage in liquid-liquid phase separation (LLPS) [3, 8-10]. Wildtype UBQLN2 accumulates in numerous neurodegenerative diseases including Huntington’s disease (HD), Lewy body dementia and amyotrophic lateral sclerosis (ALS) /frontotemporal dementia (FTD) caused by the *C9ORF72* repeat expansion[11-15]. When mutated, UBQLN2 directly causes hereditary neurodegeneration that manifests as ALS/FTD spectrum disease associated with UBQLN2 and TDP-43 accumulation [16-18].

Precisely how UBQLN2 functions in PQC pathways is not well understood, nor is it clear how UBQLN2 mutations drive neurodegeneration. In both cases, however, UBQLN2 protein-protein interactions appear to play a critical role. For example, UBQLN2 missense mutations have been shown to compromise UBQLN2 function in PQC by reducing interactions with Hsp70, limiting delivery of polyubiquitinated proteins to the proteasome, reducing autophagy, and altering liquid-liquid phase separation (LLPS) in a manner that promotes UBQLN2 accumulation and aggregation [9, 16, 19-25].

Here we sought to identify novel UBQLN2 interactors because greater knowledge of the range of UBQLN2 protein interactors will aid our understanding of how UBQLN2 acts, both in health and disease. Knowledge of the UBQLN2 interactome could also help explain why UBQLN2, principally a cytoplasmic PQC factor, translocates to the nucleus under proteotoxic stress [20]. For example, in HD mouse models, UBQLN2 localizes to neuronal nuclei where it is thought to facilitate clearance of mutant Huntingtin [13-15, 26]. Several membraneless organelles in the nucleus, notably nucleoli and PML bodies, have recently been implicated in nuclear PQC [27-30]. These subnuclear condensates undergo phase transitions from liquid-like to solid states and sequester stress-induced protein aggregates. UBQLN2 is already known to regulate the dynamics of stress granules [7], which are cytosolic condensates that modulate protein translation under conditions of stress. Conceivably, UBQLN2 plays a similar role in regulating nuclear condensates.

Here we report a novel UBQLN2 interactor, RTL8 (previously known as FAM127A or CXX1B), that readily localizes to the nucleus, regulates the subcellular localization of UBQLN2, and colocalizes with UBQLN2 in subnuclear condensates. RTL8 belongs to a family of neo-functionalized retrotransposon-derived Gag-like genes that include three highly homologous proteins, RTL8A, RTL8B and RTL8C. RTL8 proteins are largely unstudied. While no function has been ascribed to the RTL8 family, they are expressed in a wide range of human tumors [31-33]. Recently Whiteley and colleagues [34] identified RTL8 as one of several proteins whose levels are dysregulated in UBQLN2 murine disease and knockout models, consistent with our finding that RTL8 is a UBQLN2-interacting protein. Here we show that RTL8 promotes UBQLN2 localization to the nucleus, where both proteins localize to subnuclear structures that contain additional PQC components and are often adjacent to PML bodies. Moreover, RTL8 preferentially colocalizes with and promotes the nuclear localization of UBQLN2 over the two other brain-expressed ubiquilin proteins, UBQLN1 and UBQLN4. UBQLN2—more than UBQLN1 and UBQLN4— also stabilizes RTL8 levels in cells and mouse brains, pointing to functional differences among ubiquilins.

## Results

### UBQLN2 interacts with a small nuclear protein, RTL8C, that promotes UBQLN2 translocation into the nucleus

To identify novel UBQLN2-interacting proteins we performed an unbiased mass spectrometry (MS) immunoprecipitation screen using HEK293T cells transfected with FLAG-tagged WT-UBQLN2 (FLAG-UBQLN2). Repeated MS experiments detected many previously identified UBQLN2 interactors including proteasome subunits, chaperones, and other UBQLN family members (Supplementary Table 1). Among abundant interactors, we detected a 13 kDa protein, retrotransposon Gag-like protein 8C (RTL8C) (Fig. S1A). Human RTL8C (hRTL8C) is one of three members of the RTL8 family of retrotransposon-derived proteins conserved between mouse and humans, the others being RTL8A and RTL8B (Fig. 1A). Human RTL8 proteins are more than 90% homologous (Fig. S1B). Indeed, RTL8C was identified by a total of six peptides, two of which were unique to it while the other four are shared with RTL8A, leaving open the possibility that UBQLN2 interacts with more than one member of the RTL8 family. For clarity, we use the terminology RTL8 to refer to the family of proteins when antibodies are unable to differentiate them in our experiments.

**Figure 1.**
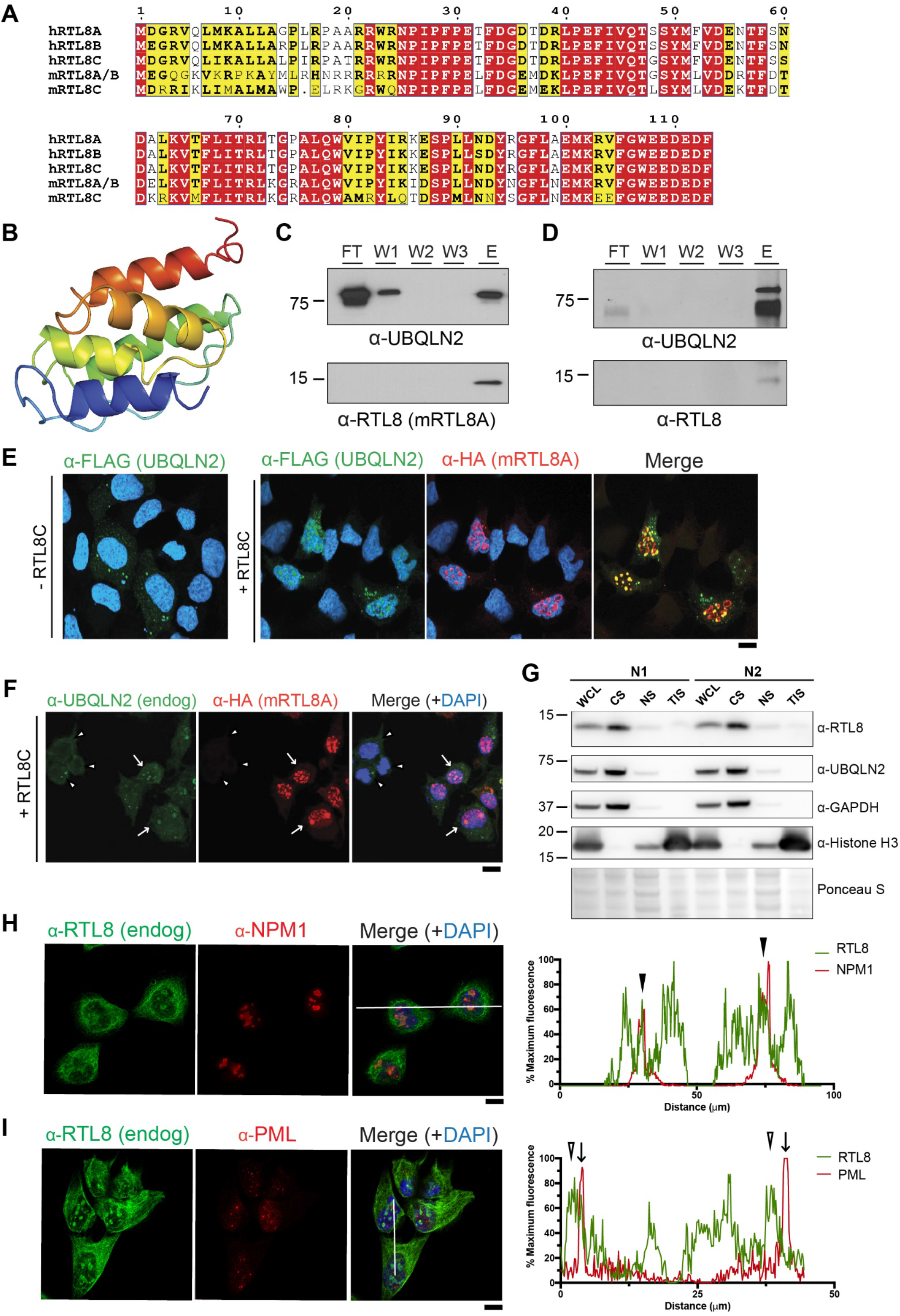
RTL8, a small family of UBQLN2 interacting proteins, promotes nuclear translocation of UBQLN2. **(A)** Alignment of the protein sequences for the three human (hRTL8) and two murine (mRTL8) RTL8 homologs, showing a high degree of sequence homology. Similar residues are boxed in yellow; the bold typeface indicates that these residues are found in the majority of RTL8 homologs. Identical residues are bolded in white font and boxed in red. **(B)** Predicted 3D structure for human RTL8C using C-I-TASSER. hRTL8C is shown in spectrum color, corresponding with blue N-terminus to red C-terminus gradient. The predicted structure consists mainly of alpha-helical regions. **(C)** Immunoblots of *in vitro* pull-down of recombinant purified protein GST-UBQLN2 and His-mRTL8A. UBQLN2-mRTL8A interaction was confirmed by detection of mRTL8A in the eluate (E), and not flow-through (FT) and washes (W1-3). **(D)** Immunoprecipitated FLAG-UBQLN2 pulled-down RTL8 from brain lysates of FLAG-tagged UBQLN2 transgenic mice. **(E)** Representative images showing mRTL8A-dependent nuclear translocation of UBQLN2 and co-localization into subnuclear structures. HEK293 cells were co-transfected with FLAG-UBQLN2 (green) and control HA-empty vector or HA-mRTL8A (left panel and right series, respectively). DAPI (blue) stained nuclei. (Scale bar = 10μm). **(F)** Representative images showing mRTL8A-mediated nuclear translocation of endogenous (endog) UBQLN2. HEK293 cells were transfected with HA-mRTL8A and stained for endogenous UBQLN2 (green) and HA-mRTL8A (red). Merged images include DAPI to highlight nuclei (blue). Arrows highlight cells demonstrating localization of endogenous UBQLN2 to nuclear puncta containing HA-mRTL8A. Arrowheads identify cells not expressing HA-mRTL8A in which UBQLN2 is diffusely cytoplasmic (Scale bar = 10μm). **(G)** Immunoblot of the subcellular distribution (WCL: whole cell lysate, CS: cytoplasmic detergent-soluble fraction, NS: nuclear detergent-soluble fraction, and TIS: total detergent-insoluble fraction) of endogenous (endog) RTL8 and UBQLN2 in HEK293 cells. GAPDH and Histone H3 served as cytoplasmic and nuclear markers, respectively. Ponceau S staining was used to normalize loading. N=3, with two technical replicates per biological replicate. **(H, I)** Representative immunofluorescence images showing co-localization of endogenous RTL8 (green) with nucleoli (H; NPM1) and PML bodies (I; PML) (red), merged with DAPI (blue). (Scale bars = 10μm). Pixel intensity plots measured across the white lines in the corresponding merged images reveal coincident peak fluorescence between RTL8 and NPM1 (indicated by closed arrow heads, H) but not in PML (arrows indicate PML peaks, open arrow heads indicate RTL8 peaks, I).

We predicted a structure model for hRTL8C using C-I-TASSER [35] and deposited it in the neXtProt database [36] at https://www.nextprot.org/entry/NX_A6ZKI3/gh/zhanglabs/COFACTOR. The predicted hRTL8C monomeric structure consists mainly of four alpha helical and loop domains (Fig. 1B) with an estimated TM-score of 0.65 (a TM-score > 0.5 suggests correct global topology [37]). This predicted structure displays homology with the capsid protein of the related retrotransposon Ty3/Gypsy [32] and *Drosophila melanogaster* retrovirus-like Arc1 protein.

To enable further *in vitro* and *in vivo* experiments, we cloned the mouse RTL8A gene (mRTL8A) which is the mouse RTL8 family member most similar to hRTL8C (Fig. S1B). To confirm a direct interaction between mRTL8A and UBQLN2, we performed pull-down experiments with recombinant GST-tagged UBQLN2 and His-tagged mRTL8A. GST-UBQLN2 purified protein was applied to a GST agarose column to which His-mRTL8A was then added. When reduced glutathione was used to elute UBQLN2, immunoblot analysis revealed mRTL8A in the eluate, confirming a direct interaction between the two proteins (Fig. 1C).

Confirmation of interaction between UBQLN2 and endogenous RTL8 was obtained in mouse brain by immunoprecipitating FLAG-UBQLN2 from transgenic mice expressing a low level of wildtype FLAG-UBQLN2 [22] Mouse brain lysate from UBQLN2 transgenic mice was incubated with anti-FLAG agarose beads and, after washing, FLAG-UBQLN2 was eluted from the beads. Immunoblot analysis demonstrated the presence of RTL8 in the eluate (Fig. 1D), confirming an *in vivo* interaction between the two proteins in brain.

To determine whether UBQLN2 and mRTL8A interact and co-localize *in vivo*, HEK293 cells were co-transfected with FLAG-UBQLN2 and HA-tagged mRTL8A (HA-mRTL8A). Expressed alone, FLAG-UBQLN2 was found diffusely and in small puncta in the cytoplasm (Fig. 1E, left panel) as described previously [20, 22]. When co-transfected with HA-mRTL8A, however, FLAG-UBQLN2 translocated into the nucleus where it co-localized with mRTL8A (Fig. 1E, right panel). HEK293 cells transfected with HA-mRTL8A alone recruited endogenous UBQLN2 into the nucleus (Fig. 1F, white arrows), whereas endogenous UBQLN2 remained largely cytoplasmic in untransfected cells (Fig. 1F, white arrowheads).

To determine the subcellular localization of RTL8, we performed biochemical fractionation on HEK293 cells. We probed whole cell lysate (WCL), detergent-soluble cytoplasmic (CS) and nuclear fractions (NF), and total detergent-insoluble fraction (TIS) by immunoblot with a commercially available RTL8 antibody (Fig. 1G). RTL8 proteins mostly resided in the cytoplasmic fraction but were also found in the nuclear and detergent-insoluble fractions. UBQLN2 was present only in the detergent-soluble fractions and mostly in the cytoplasmic fraction.

Overexpressed HA-mRTL8A concentrates in nuclear puncta (Fig. 1F, middle panel) suggesting that it partitions into specific subnuclear organelles. The co-localization with UBQLN2, a known participant in protein degradation pathways, led us to investigate whether endogenous RTL8 resides in nucleoli or PML nuclear bodies, both of which have been implicated in nuclear PQC [27-30]. Unfortunately, commercial α-RTL8 antibodies do not stain endogenous RTL8 in HEK293 cells by immunofluorescence, thus we generated an α-RTL8A polyclonal rabbit antibody. This antibody detected endogenous nuclear RTL8 staining in subnuclear structures that morphologically resemble those seen when HA-tagged mRTL8A is overexpressed (Fig. 1E and 1F). Probing with antibodies for nucleoli (Fig. 1H) or PML nuclear bodies (Fig. 1I) revealed robust localization of RTL8 to nucleoli exclusively.

### UBQLN2 redistributes mRTL8A within sites of nuclear PQC

UBQLN2 and mRTL8A co-localize to distinct puncta in the nucleus, as shown above. We sought to determine whether, as seen with endogenous RTL8, these structures represent nucleoli. To do so, we transiently expressed mRTL8A in HEK293 cells with or without FLAG-UBQLN2. We noted intriguing changes in the subnuclear localization of mRTL8A when UBQLN2 was co-expressed. Expressed alone, mRTL8A tended to co-localize with the nucleolar protein NPM1, mirroring endogenous RTL8 staining (Figs. 2A and 2C (*-UBQLN2*)). In contrast to endogenous RTL8, however, overexpressed mRTL8A concentrates in close proximity to PML bodies (Figs. 2B and 2E (*-UBQLN2*)). Interestingly, UBQLN2 on its own did not colocalize with either NPM1 or PML (Fig. S2A). Rather, in cells expressing both UBQLN2 and mRTL8A, we see two populations of mRTL8A puncta: those that are colocalized with UBQLN2 (UBQLN2/mRTL8A puncta), and those that are not.

**Figure 2:**
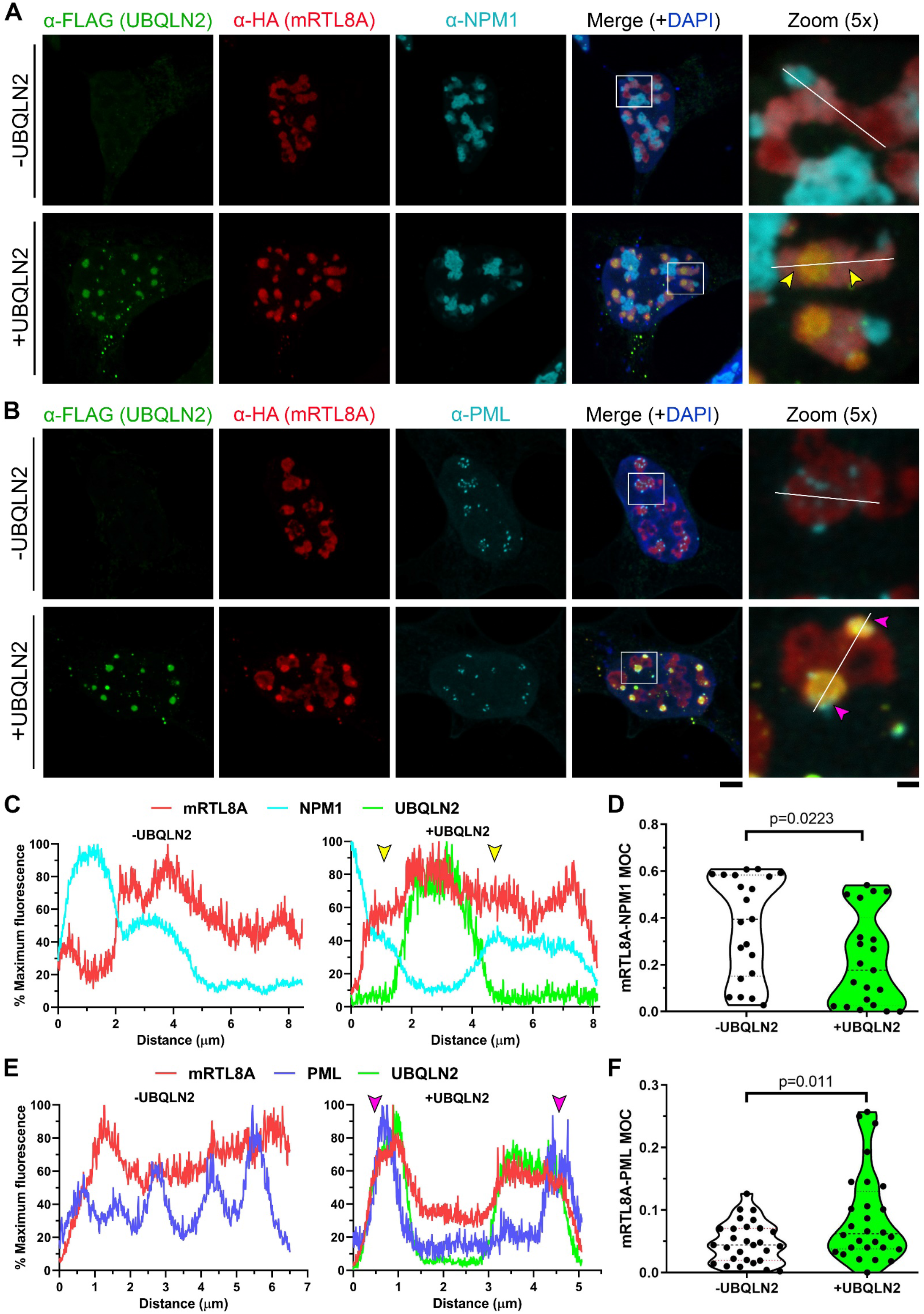
UBQLN2-mRTL8A subnuclear structures are distinct from nucleoli and recruit PML bodies. **(A), (B)** Representative images of HEK293 cells co-transfected with HA-mRTL8A (*red*) and either an empty vector (-UBQLN2) or one encoding FLAG-UBQLN2 (+UBQLN2) (*green*). Cells were stained for **(A)** NPM1 or **(B)** all PML isoforms (*cyan*). (Scale bar for merged images = 5 µm and zoomed images = 1 µm). Yellow arrows in (A) and (C) represent regions surrounding UBQLN2-mRTL8A puncta that co-stain for NPM1 and mRTL8A, but exclude UBQLN2. Magenta arrows in (B) and (E) represent regions surrounding UBQLN2-mRTL8A puncta that are in close proximity with multiple PML bodies. **(C)** Pixel intensity plots measured across the white lines in the merged 5x magnification images in (A). Raw values were normalized to the maximum fluorescence value for each channel and plotted as a function of distance. **(D)** Quantification of co-localization of mRTL8A with NPM1 via the Mander’s overlap coefficient (MOC) in the presence and absence of overexpressed UBQLN2. Outliers as determined by a Grubbs’ test were excluded following which an unpaired two-tailed Student’s t-test was used to test for statistical significance. N=21 cells from 2 biological replicates. **(E)** Pixel intensity plots measured across the white lines in the merged 5x magnification images in (B). Raw values were normalized to the maximum fluorescence value for each channel and plotted as a function of distance. **(F)** Quantification of co-localization of mRTL8A with PML via MOC in the presence and absence of overexpressed UBQLN2. Outliers as determined by a Grubbs’ test were excluded, following which an unpaired two-tailed Student’s t-test was used to test for statistical significance. N=28 cells from 3 biological replicates.

The mRTL8A puncta that exclude UBLQN2 co-stain with NPM1, suggesting colocalization with nucleoli, whereas UBQLN2/mRTL8A puncta are not positive for NPM1 (Figs. 2A and 2C (*+UBQLN2, bottom panels*); and Fig. S2B). Interestingly, UBQLN2/mRTL8A puncta frequently show heterogeneity, with some mRTL8A signal not overlapping with UBQLN2 staining. The mRTL8A signal that excludes UBQLN2 is often positive for NPM1 and seen as a “ring” of NPM1 signal that surrounds, but does not overlap, the UBQLN2 signal (yellow arrows in Figs. 2A and 2C (*+UBQLN2*)). Quantification of the Mander’s overlap coefficient revealed decreased colocalization of mRTL8A with NPM1 when UBQLN2 was coexpressed (Fig. 2D), suggesting that UBQLN2 sequesters mRTL8A away from nucleoli and into UBQLN2/mRTL8A puncta.

PML bodies were also observed in close proximity to UBQLN2/mRTL8A puncta, with multiple PML bodies preferentially colocalizing with and clustering around these puncta (magenta arrows in Figs. 2B and 2E (-*+UBQLN2*); and Fig. S2C). Quantification of the Mander’s overlap coefficient revealed increased colocalization of mRTL8A with PML bodies when UBQLN2 was coexpressed (Fig 2E). Taken together, these data indicate that UBQLN2 promotes redistribution of mRTL8A away from nucleoli and into close proximity to PML bodies.

### mRTL8A co-localizes with nuclear PQC components, both in the presence and absence of UBQLN2

The above results show that mRTL8A puncta colocalize with nucleoli and are closely aligned with PML bodies, both of which have been implicated in PQC. To investigate the properties of mRTL8A puncta we immunostained transfected cells for the endogenous PQC markers Hsp70, SQSTM1/p62 and ubiquitin, as well as for the disease-associated RNA-binding protein TDP-43. Hsp70 proteins are cytoplasmic chaperones that translocate to nucleoli during proteotoxic stress and aid in protein refolding [38, 39]. SQSTM1/p62 is an autophagy receptor that facilitates nuclear PQC by translocating to the nucleus and engaging polyubiquitinated substrates [40], and is recruited to nuclear aggregates in various neurodegenerative diseases [41, 42]. TDP-43 is known to shuttle between the nucleus and cytoplasm, accumulate in the cytoplasm in several neurodegenerative diseases [43], and form nuclear aggregates with certain proteotoxic stressors [44].

Assessed by immunofluorescence, mRTL8A puncta co-localized extensively with Hsp70 both in the presence and absence of UBQLN2 (Figs. 3A and S3B). Hsp70 colocalization was greatest in intensely stained mRTL8A-positive puncta but was also seen in lower-intensity mRTL8 puncta. Overall Hsp70 fluorescence signal tended to be higher in mRTL8A-expressing cells than in neighboring untransfected cells. Whereas Hsp70 staining was seen in both high and low intensity mRTL8 puncta, p62 and ubiquitin co-localized only to the higher intensity mRTL8A puncta (Figs. 3B and 3C) that do not possess nucleolar morphology. Overexpressed UBQLN2 also tended to co-localize to high intensity mRTL8A puncta. In contrast, mRTL8A did not colocalize with TDP-43 whether in the presence or absence of UBQLN2 (Fig. 3D). UBQLN2 on its own (i.e. without overexpressed mRTL8A) colocalized only with p62 and ubiquitin (Fig. S3A-D). Interestingly, infrequently observed UBQLN2-mRTL8A cytoplasmic puncta do not co-stain with Hsp70 or ubiquitin, implying subcellular specificity with respect to recruitment of PQC components.

**Figure 3.**
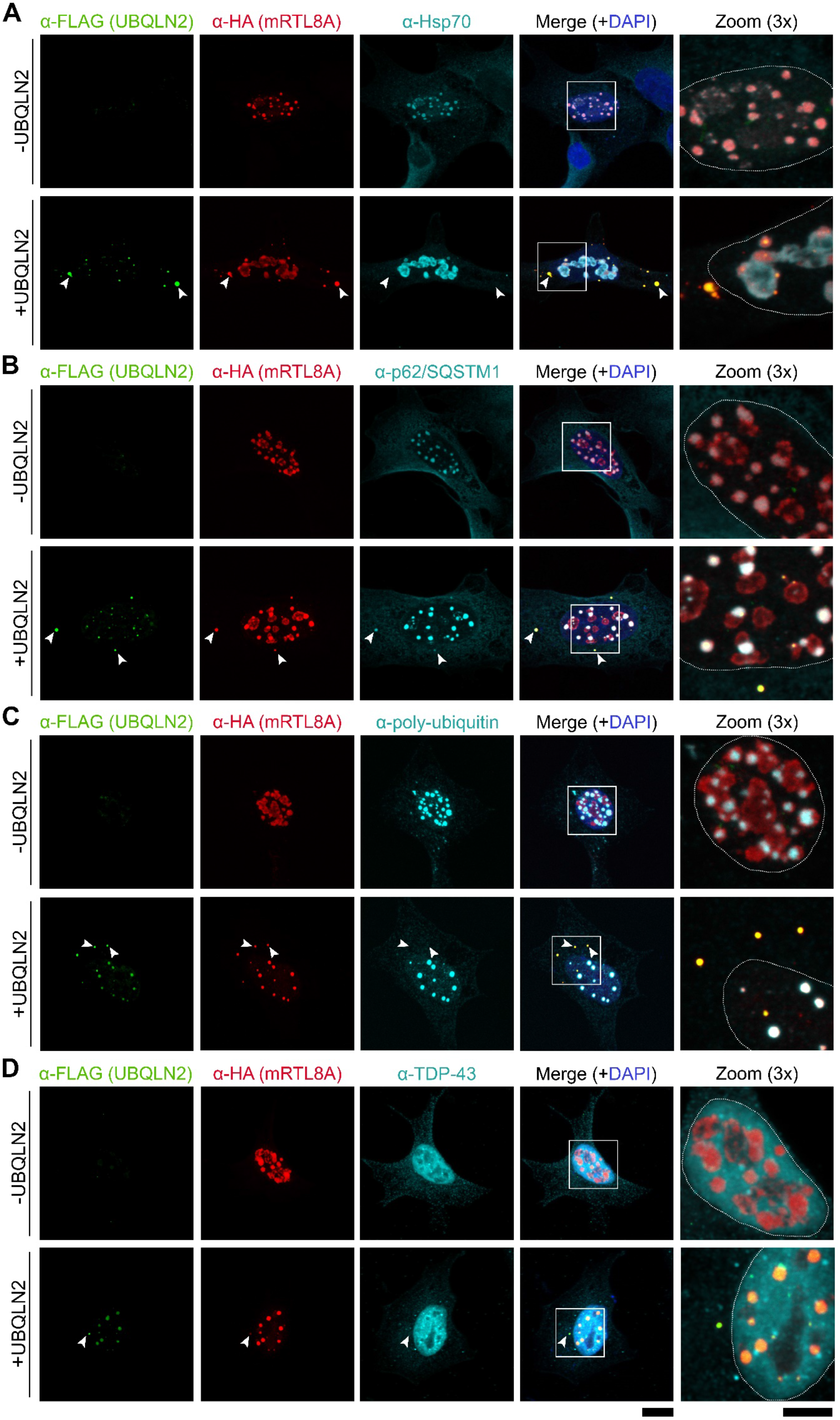
UBQLN2-mRTL8A subnuclear structures co-localize with Hsp70, p62 and ubiquitin but not TDP-43. Representative images of HEK293 cells co-transfected with HA-mRTL8A (*red*) and either empty vector or FLAG-UBQLN2 (*green*), and stained for **(A)** Hsp70, **(B)** p62, **(C)** poly-ubiquitin or **(D)** TDP-43 (*cyan*). Arrows indicate infrequent cytoplasmic UBQLN2-mRTL8A structures that are positive only for p62. White outlines in zoomed images represent the nucleus as determined by DAPI staining (blue in merged images). Figure S2E shows Hsp70 staining of cells expressing lower levels of FLAG-UBQLN2 and HA-mRTL8A compared to Figure 2A. (Scale bar for merged images = 10 µm and zoomed images = 5 µm).

In summary, overexpressed mRTL8A concentrates in two discrete subnuclear structures containing different combinations of PQC markers. The first resemble nucleoli and co-stain only for Hsp70 while the second are intensely mRTL8A-positive puncta that co-localize with Hsp70, p62 and ubiquitin. When coexpressed with mRTL8A, UBQLN2 is recruited to the latter structures.

### UBQLN2’s UBA and UBL domains are not essential for the UBQLN2-mRTL8A interaction

To begin defining the structural requirements for UBQLN2’s interaction with RTL8 we deleted two key domains of UBQLN: the UBL and UBA domains (Fig. 4A). The N-terminal UBL domain facilitates interaction with the 26S proteasome while the C-terminal UBA domain allows binding to polyubiquitinated substrates [1, 3, 8, 45, 46]. In pull-down experiments with recombinant UBQLN2 lacking either the UBA (ΔUBA) or the UBL (ΔUBL) (Fig. 4B), similar levels of mRTL8A were pulled down by both deletion constructs as shown earlier for full-length UBQLN2 (Fig. 1C). Thus, neither the UBA nor the UBL domain is essential for UBQLN2 interaction with RTL8.

**Figure 4.**
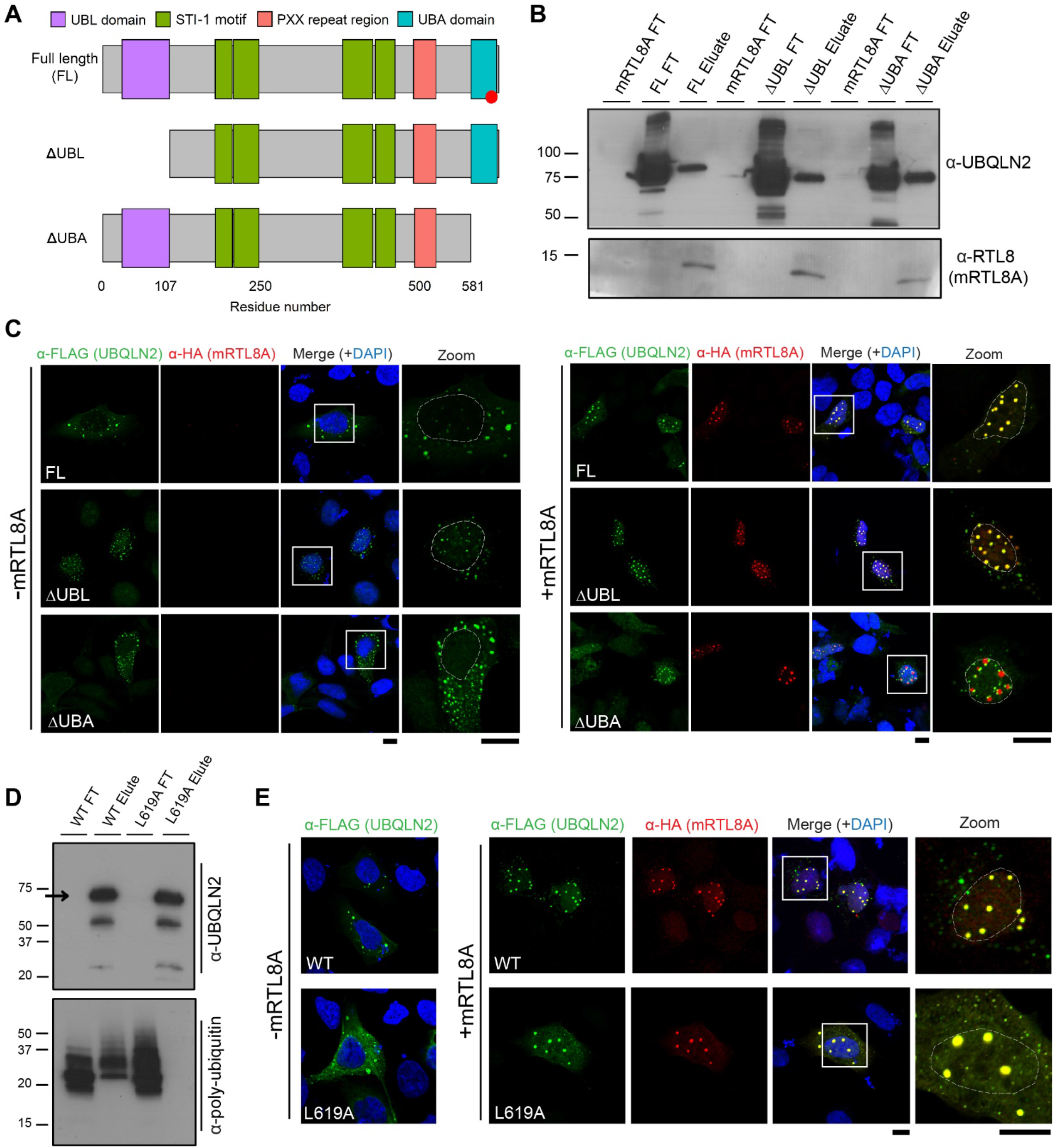
UBQLN2-mRTL8A interaction occurs independent of UBA, UBL, and ubiquitin binding domains. **(A)** Schematic depicting UBQLN2 domain deletion constructs lacking either the UBL (ΔUBL) or UBA (ΔUBA) domains, and the ubiquitin-binding deficient L619A mutation (represented as a red circle). **(B)** Pull-down of recombinant His-mRTL8A by recombinant full-length GST-UBQLN2 full-length (FL), ΔUBL, and ΔUBA proteins. The flow-through (FT) and eluate were visualized on an immunoblot with α-UBQLN2 (top) or α-RTL8 antibodies (bottom). **(C)** Representative images of transfected HEK293 cells expressing FLAG-UBQLN2 WT, ΔUBA or ΔUBL (green) without (left panel) or with (right panel) HA-mRTL8A *(red)*. White outlines in zoomed images delimit the nucleus as determined by DAPI staining (blue in merged images). (Scale bars = 10 µm). **(D)** Pull-down experiment confirming the loss of poly-ubiquitin binding by recombinant GST-UBQLN2 L619A protein. The flow-through (FT) and eluate were visualized by immunoblot with α-UBQLN2 (top) or α-poly-ubiquitin antibody (bottom). *Arrow* denotes UBQLN2. **(E)** Representative images showed continued co-localization and nuclear localization of mutant UBQLN2 L619A with mRTL8A. HEK293 cells were transfected to express FLAG-UBQLN2 WT (top) or L619A (bottom) (green) without (*left*) or with (*right*) HA-mRTL8A (red). White outlines in zoomed images delimit the nucleus determined by DAPI staining (blue in merged images). (Scale bars = 10 µm).

Next, we tested whether deleting either domain affected the ability of UBQLN2 to co-localize with mRTL8A and translocate to the nucleus. In the absence of co-expressed mRTL8A, cells expressing full-length UBQLN2 or UBQLN2 deletion constructs showed a similar distribution, predominantly in the cytosol with faint speckle-like staining in the nucleus (Fig. 4C, *left*). When co-expressed with mRTL8A, ΔUBL-UBQLN2 behaved identically to full-length UBQLN2 in translocating to the nucleus and colocalizing with mRTL8A in subnuclear structures (Fig. 4C, *right*). In contrast, ΔUBA-UBQLN2 only partially co-localized with mRTL8A even though its nuclear localization was still promoted by mRTL8A. Puncta of mRTL8A and ΔUBA-UBQLN2 also differed in that they were juxtaposed rather than fully colocalized, as seen with full-length and ΔUBL-UBQLN2.

The above results suggest that the UBA domain plays a significant role in the interaction between RTL8 and UBQLN2. Since UBA is the polyubiquitin binding domain of UBQLN2, we tested whether ubiquitin binding itself modulates the UBQLN2-RTL8 interaction. Residue L619 in the UBA domain is essential for polyubiquitin binding by UBQLN2 [20, 22], which we verified in polyubiquitin pulldown experiments with UBQLN2 mutated at this residue (L619A, Fig. 4D). In transfected HEK293 cells, both WT and L619A UBQLN2 are predominantly cytoplasmic (Fig. 4E). When co-expressed with mRTL8A, L619A UBQLN2 co-localized with mRTL8A in the nucleus, nearly identically to wild-type UBQLN2. This result implies that polyubiquitin binding is not essential for UBQLN2 to engage mRTL8A and that other features of the UBA domain, beyond its ability to bind ubiquitin, modulate the interaction with RTL8. Although neither the UBA or UBL domain nor an ability to bind ubiquitin chains is required for UBQLN2 to interact with mRTL8A, deletion of the UBA domain may elicit structural changes in UBQLN2 that alter nuclear colocalization with mRTL8A.

### Preferential interactions of RTL8 with UBQLN2 support functional heterogeneity among ubiquilins

Other UBQLN family members, including UBQLN1 and UBQLN4, share conserved structural domains with UBQLN2 except for the proline-rich repeat region (PXX) that is unique to UBQLN2 [47] (Fig. 5A). This similarity prompted us to test whether mRTL8A also co-localizes with or aids UBQLN1 and UBQLN4 translocation to the nucleus. When expressed alone in HEK293 cells, UBQLN1 and UBQLN4 exhibited subcellular distributions distinct from UBQLN2 (Fig. 5B, *upper panels*). Whereas UBQLN2 was present mostly in cytoplasmic puncta with rare speckle-like nuclear staining, both UBQLN1 and UBQLN4 showed both diffuse cytoplasmic and nuclear staining. When individual UBQLNs were overexpressed with mRTL8A, UBQLN2 usually co-localized with mRTL8A, primarily in the nucleus, whereas UBQLN1 showed highly variable co-localization (Fig. 5B, *lower panels*) and UBQLN4 did not co-localize with mRTL8A. Using Mander’s overlap coefficient to quantify co-localization, we confirmed significant differences in the degree of mRTL8A co-localization with UBQLNs 1, 2 and 4 (Fig. 5C). We also quantified the nuclear to cytoplasmic (N/C) ratio of all three UBQLNs (Fig. 5D). In the absence of exogenous mRTL8A, UBQLN2 had the lowest N/C ratio, with the two other UBQLNs having N/C ratios closer to one. Co-expression of mRTL8A led to an almost two-fold increase in the N/C ratio of UBQLN2 with no appreciable effect on the N/C ratio of UBQLN1 or UBQLN4.

**Figure 5.**
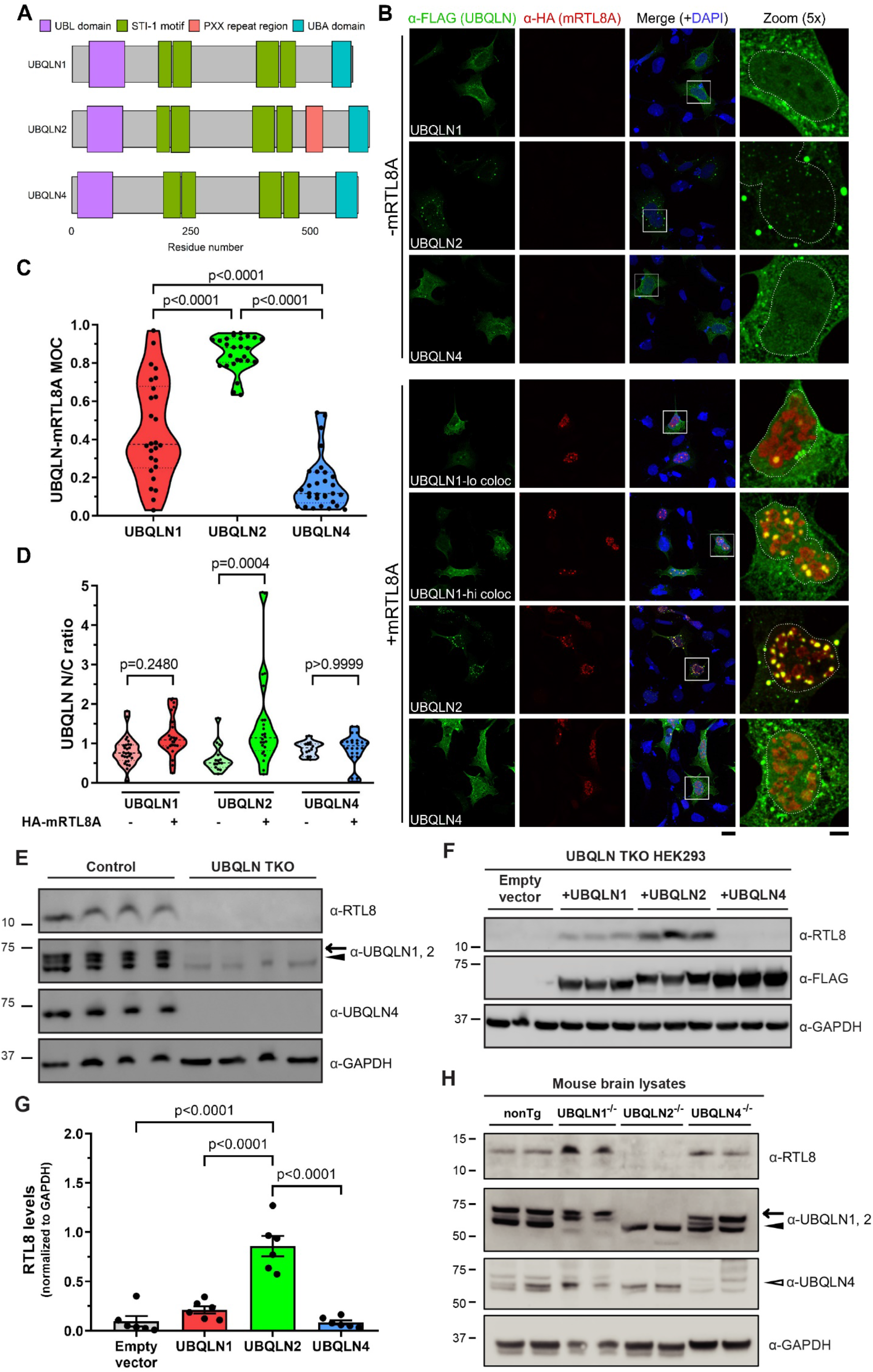
Among brain-expressed ubiquilins, UBQLN2 preferentially colocalizes with mRTL8A and regulates endogenous RTL8 levels in cells and mouse brain tissue. **(A)** Schematic of the three brain-expressed ubiquilin proteins (UBQLNs 1,2 and 4) showing common structural motifs and the PXX repeat region that distinguishes UBQLN2. **(B)** Representative immunofluorescence images of HEK293 cells co-transfected with HA-mRTL8A (red) and either FLAG-tagged UBQLN1, UBQLN2 or UBQLN4 (green), shown merged with DAPI (blue). White outlines in the zoomed images delimit the nucleus, determined by DAPI staining. Representative images of high and low co-localization of UBQLN1 with mRTL8A are included to illustrate the wide range of observed co-localization for UBQLN1 (lo coloc= low co-localization, hi coloc = high co-localization). (Scale bar for merged images = 20 µm and zoomed images= 5 µm). **(C)** Quantification of co-localization of mRTL8A with UBQLN1, UBQLN2 or UBQLN4. Mander’s correlation coefficient (MOC) was calculated for each cell expressing both constructs in a field of view. Outliers as determined by a Grubbs’ test were excluded, following which a one-way ANOVA (p<0.0001) and a Tukey’s multiple comparisons test were performed to identify statistically significant differences among Mander’s correlation coefficients. 26, 27 and 31 cells were analyzed for UBQLN1, UBQLN2 and UBQLN4, respectively, from 2 biological replicates. **(D)** Quantification of UBQLN nuclear localization, represented by the ratio of nuclear to cytoplasmic (N/C) fluorescence intensity. Statistical significance was determined by first excluding outliers as determined by a Grubbs’ test and then performing a one-way ANOVA (p <0.0001) followed by a Tukey’s multiple comparisons test. 28, 15 and 14 cells were analyzed for UBQLN1, UBQLN2 and UBQLN4, respectively, without mRTL8A. 25, 24 and 24 cells were analyzed for UBQLN1, UBQLN2 and UBQLN4, respectively, with mRTL8A. 2 biological replicates were used in both cases. **(E)** Immunoblot of lysates from HEK293 T-rex control (containing UBQLN1, 2 and 4) and UBQLN1, 2 and 4 total knockout (TKO) HEK293 cells visualizing the loss of endogenous RTL8 proteins in TKO cells. Arrow and arrowhead denote bands corresponding to UBQLN2 and UBQLN1, respectively. GAPDH was used as a loading control. **(F)** Representative immunoblots for endogenous RTL8 levels in TKO HEK293 cells transfected either with an empty vector, FLAG-tagged UBQLN1, UBQLN2 or UBQLN4. GAPDH was used as a loading control. **(G)** Relative levels of RTL8 normalized to GAPDH quantified by densitometry. Statistical significance as determined by a one-way ANOVA (p<0.0001) followed by a Tukey’s multiple comparisons test. Data shown represent means +/-SEM normalized to UBQLN2. N=6 per condition. **(H)** Immunoblot of brain lysates from non-transgenic (nonTg), UBQLN1^-/-^, UBQLN2^-/-^ and UBQLN4^-/-^ mice. Arrows indicate each UBQLN paralog. GAPDH was used as a loading control.

A recent study demonstrated that HEK293 cells with all three UBQLNs knocked out (TKO) have no detectable RTL8 protein levels [34]. We confirmed the loss of RTL8 protein expression in TKO cells (Fig. 5E). We then assessed whether expressing UBQLN1, UBQLN2 or UBQLN4 separately could restore RTL8 protein levels in TKO cells. Based on Fig. 5B, we hypothesized that UBQLN2 and possibly UBQLN1 would rescue RTL8 expression whereas UBQLN4 would not. Indeed, UBQLN2 rescued RTL8 levels to the greatest extent while UBQLN1 slightly rescued RTL8 levels and UBQLN4 had no effect (Figs. 5F and 5G).

To determine if this differential effect of UBQLNs on RTL8 expression held true in a more physiologically relevant model, we examined RTL8 levels in brain lysates from UBQLN1, UBQLN2 and UBQLN4 knockout mice. While UBQLN2 knockout mice showed no detectable RTL8 protein, both UBQLN1 and UBQLN4 knockout mice showed RTL8 levels similar to that of WT non-transgenic (nonTg) mice (Fig. 5H). Thus, in mouse brain neither UBQLN1 nor UBQLN4 can compensate for the loss of UBQLN2. Conversely, endogenous UBQLN2 expression is sufficient to maintain RTL8 levels in UBQLN1 or UBQLN4 knockout mice. Together, these data suggest that the functional interaction between RTL8 and UBQLN2 does not extend to other brain-expressed UBQLNs.

## Discussion

UBQLN2 is known to participate in cytoplasmic PQC through the UPS and autophagy. It is less clear what role UBQLN2 plays in the nucleus, to which it translocates under various stress conditions [12, 13, 15, 20]. Nuclear aggregates of disease proteins, including known UBQLN2 substrates, are common in various neurodegenerative diseases. Previous work in yeast has demonstrated that the UBQLN2 ortholog, Dsk2, is specifically required for clearance of nuclear misfolded proteins [48].Conceivably, the recruitment of UBQLN2 to the nucleus facilitates the degradation of misfolded proteins associated with neurodegenerative proteinopathies.

While UBQLN4 reportedly functions in DNA double-stranded break repair [49], a common nuclear function across the ubiquilin family is unknown. Here, we identified a novel UBQLN2-interacting protein, RTL8, that regulates UBQLN2’s translocation to and behavior in the nucleus, and underscores functional heterogeneity within the UBQLN protein family.

RTL8 is a small protein of unknown function expressed in the cytoplasm and nucleus. Endogenous nuclear RTL8 localizes to nucleoli, which are condensates associated with ribosomal biogenesis and protein quality control [28, 50, 51]. When overexpressed, mRTL8A tends to concentrate in the nucleus and continues to co-localize to nucleoli as well as to ubiquitin-enriched subnuclear structures distinct from nucleoli. Overexpressed mRTL8A also recruits and sequesters UBQLN2 within nuclear condensates that contain other PQC proteins. Among brain-expressed ubiquilins, only UBQLN2 shows mRTL8A-dependent nuclear recruitment and colocalization: by contrast, mRTL8A only rarely or occasionally colocalizes with UBQLN4 and UBQLN1, respectively, and fails to promote nuclear translocation of either. Furthermore, UBQLN2 stabilizes levels of RTL8 protein in mouse brain, which is not the case for UBQLN1 and UBQLN4. Of the three brain-expressed UBQLNs, only the loss of UBQLN2 reduces RTL8 levels, supporting a physiological role for the interaction between these two proteins. These findings suggest that RTL8 preferentially regulates UBQLN2 function in the nucleus over UBQLN1 or UBQLN4.

Limitations to this study are the small number of replicates in our MS screen identifying RTL8 as a potential interactor and the need to employ overexpression in our analysis of RTL8 because of the lack of specific antibodies suitable for detecting the endogenous protein. We were nonetheless able to validate the direct interaction between mRTL8A and UBQLN2 through recombinant protein pull-downs and immunoprecipitation from mouse brain lysate. Given the approximately 7-fold enrichment of RTL8 in our study, it is notable that earlier MS screens for UBQLN2 interactors did not identify RTL8 [7, 20]. One possible explanation for this discrepancy is that our MS screen was performed in HEK293 cells which contain multiple copies of the q arm of the X chromosome, the locus of the RTL8 gene family. Thus, HEK293 cells likely express higher levels of RTL8 than other cell models or tissues, enabling detection in our screen. Recent literature also supports our findings: A proteomics study examining the effects of mutations or knockout of UBQLN2 identified RTL8 as one of several gag/pol proteins that are highly regulated by UBQLN2 [34].

Multiple structural motifs in UBQLN2 could mediate the direct interaction with RTL8 including the ubiquitin-like domain (UBL), the ubiquitin-associated domain (UBA), the four STI1 motifs, and the proline-rich repeat (PXX) region. Our recombinant protein pull-down and immunofluorescence experiments establish that mRTL8A functionally interacts with UBQLN2 independent of the UBA or UBL domains and does not depend on ubiquitin binding. These findings suggest regions outside these domains regulate the interaction of RTL8 and UBQLN2. The preferential interaction of mRTL8A with UBQLN2 over UBQLN1 and UBQLN4, points to the distinctive PXX domain of UBQLN2 as a potentially critical motif. At the same time, some of our results imply that the functional interaction between UBQLN2 and mRTL8A is not regulated by a single domain in UBQLN2. First, overexpressed UBQLN1 also partially colocalizes with and rescues RTL8 levels in UBQLN knockout cells, albeit much less so than UBQLN2. While this finding contrasts with our finding that UBQLN1 does not stabilize RTL8 levels in mouse brain, we hypothesize that overexpressed UBQLN1 engages in a lower-affinity interaction with RTL8. Second, overexpressed UBQLN2 lacking the UBA domain co-localizes with mRTL8A less robustly then full-length UBQLN2, suggesting the interaction with RTL8 is partially mediated by the UBA domain. UBQLN1 and UBQLN2 are 96% identical in the UBA domain whereas UBQLN4 and UBQLN2 are only 88% identical in this domain. If the UBA domain is an important regulator of UBQLN-RTL8 interaction, the greater similarity of UBQLN1 than UBQLN4 to UBQLN2 in this domain may explain the partial functional interaction of UBQLN1 with RTL8.

Identifying the motifs that determine RTL8 interaction with UBQLN2 will offer insight into how RTL8 regulates UBQLN2 function in health and dysfunction in disease. For example, only UBQLN2 contains the PXX domain where the majority of fALS-causing mutations accumulate. If the PXX domain is important for RTL8 binding this could have important disease implications. fALS mutations in UBQLN2 that alter RTL8 binding may directly or indirectly impact disease. Likewise, involvement of the UBA domain in RTL8 binding would have implications for UBQLN2 function. Although ubiquitin binding to the UBA domain appears not to be required for RTL8 binding to and nuclear recruitment of UBQLN2, RTL8 binding to UBQLN2 could modulate ubiquitin binding by UBQLN2. Intriguingly, all three UBQLNs undergo liquid-liquid-phase separation and are inherently aggregate-prone [52], and the UBA domain itself may seed UBQLN oligomerization and aggregation [22]. A systematic study of structural motifs and disease mutations in UBQLN2 will be required to determine which regions are critical for binding and whether fALS mutations impact RTL8 binding or RTL8-dependent properties of UBQLN2.

The function of RTL8 and its family members is unknown, but the data reported here suggest potential roles for RTL8. Both endogenous RTL8 and overexpressed mRTL8A localize to the nucleolus, a nuclear organelle recently implicated in PQC [28]. Overexpressed mRTL8A colocalizes with the chaperone Hsp70 in nucleoli as well. When co-expressed, mRTL8A and UBQLN2 colocalize in puncta that recruit PQC markers. Our data are consistent with RTL8 contributing to PQC pathways in the nucleus. Interestingly, endogenous RTL8 resides in both the nucleus and cytoplasm, but when overexpressed, mRTL8A is found almost entirely within the nucleus. Many proteins, including UBQLN2 and Hsp70, translocate to the nucleus under stress conditions. RTL8 may be similarly regulated by stress and, upon movement to the nucleus, recruit additional PQC proteins as well.

RTL8 puncta colocalize with nucleolar structures and are closely associated with PML bodies. PML bodies perform diverse functions in the nucleus, including serving as sites of proteasomal degradation [29, 53, 54]. As opposed to nucleoli, which are constitutive PQC compartments [28], PML bodies are thought to serve as stress inducible PQC compartments [29]. Substrate proteins targeted to PML bodies tend to be SUMOylated, with the PML protein itself acting as a SUMO E3 ligase [54]. There is extensive interconnectivity between SUMOylation and ubiquitination pathways [55-57], highlighting a potential role for UBQLN2 in shuttling substrates that may be SUMOylated and/or ubiquitinated. Since mRTL8A resides in close proximity to PML bodies, it may facilitate recruitment of UBQLN2 to this nuclear site of proteasomal degradation. Alternatively, the PML clustering we observe could represent a byproduct of protein aggregation rather than a specific functional role for mRTL8A [53, 58]. RTL8 proteins, including mRTL8A, have a predicted SUMOylation site (Lys85) which may target them for degradation (http://jassa.fr/) [59]. We do not think that PML clustering is an artifact of mRTL8A aggregation, however, based on unpublished control studies which assessed PML clustering on mutant TDP43 which forms nuclear aggregates [60] and is itself a target for SUMOylation [61, 62]. This data suggests that the PML clustering on mRTL8A puncta is more complex than PML-mediated degradation of aggregated proteins. Further experiments will be required to clarify the significance of this observed PML clustering.

UBQLN2 has been implicated in a wide range of neurodegenerative diseases, with nuclear aggregates of misfolded proteins representing a signature pathological feature for several of them [12, 13, 15, 20]. Understanding the role that UBQLN2 plays in nuclear PQC and the mechanisms and critical components that regulate it will require further study. The data presented here show that RTL8, a novel protein that binds UBQLN2, drives UBQLN2 into the nucleus where they form condensates that recruit PQC components. This interaction with and functional effect on UBQLN2 is selective for this particular UBQLN and does not occur with UBQLN1 and UBQLN4. Future investigation into the significance of the RTL8/UBQLN2 interaction and whether it is affected by known human disease mutations may help decipher the mechanisms by which UBQLN2 dysfunction leads to neurodegenerative disease.

## Materials and methods

### Mass spectrometry (MS) and data analysis

UBQLN2 complexes were isolated from HEK293 cells overexpressing FLAG-tagged wild-type UBQLN2 using anti-FLAG M2 Affinity Gel (Millipore-Sigma, A2220). After washing, the gel beads were eluted with FLAG peptide and the eluates were analyzed at the University of Michigan Mass Spectrometry-Based Proteomics Resource Facility (https://www.pathology.med.umich.edu/proteomics-resource-facility).

Upon cysteine reduction (10 mM DTT) and alkylation (65 mM 2-chloroacetamide or iodoacetamide, with similar results) of the cysteines, the isolated proteins were digested overnight with sequencing grade modified trypsin (Promega). The resulting peptides were resolved on a nano-capillary reverse phase column (PicoFrit column, New Objective) using a 1% acetic acid/acetonitrile gradient at 300 nl/min and directly introduced into a linear ion-trap mass spectrometer (LTQ Orbitrap XL, Thermo Fisher). Data-dependent MS/MS spectra on the five most intense ions from each full MS scan were collected (relative collision energy ∼35%).

Proteins and peptides were identified by searching the data against *H. sapiens* protein database containing only the canonical, reviewed protein entries (Swiss-Prot, 20286 entries; downloads release 2020_05) using Proteome Discoverer (v2.4, ThermoFisher Scientific). Enzyme specificity was set to fully tryptic digestion with up to 2 missed cleavages. Search parameters included a precursor and fragment mass tolerances of 50 ppm and 0.6 Da, respectively. Carbamidomethylation of cysteine was considered fixed modification. Oxidation of methionine, deamidation of asparagine and glutamine, and ubiquitination (diglycine remnant) on lysine residues were considered as variable modifications. Percolator, a peptide-to-spectrum (PSM) validator, was used to filter the protein/peptides to retain only those with a false discovery rate (FDR) of ≤1%.

To correct for missing/zero PSM values in control samples since natural log of zero leads to a numerical error, an empirical value of 1 was added to PSM values of both control and FLAG-UBQLN2 datasets. Data analysis was done on RStudio (v 1.2.5033) using the normalized spectrum abundance factor (NSAF) method outlined by Zybailov and colleagues [63]. Each protein’s PSM value was first normalized to its length, denoted as length normalized PSM. Next, the length-normalized PSM value for each protein was divided by the sum of all length normalized PSMs for the given sample, denoted as NSAF. To calculate fold enrichment over control, NSAF values for each hit from a FLAG-UBQLN2 dataset were normalized to their respective control PSM values. These values were then log_2_ transformed and tested for significance using a Student’s t-test. Volcano plot was generated using the ggplot2 package [64] by plotting log_2_ transformed fold enrichment values and negative log_10_ transformed p-values on the x and y axes respectively. The p-value threshold was expanded to 0.15 (-log_10_ value of 0.8239) to prevent the exclusion of possible interactors since only two replicates were used for this analysis. Hits with a log_2_ transformed fold enrichment of greater than 1.5 and a p-value of 0.15 (-log_10_ value of 0.8239) were considered possible interactors, and are shown in red.

### Antibodies

The antibodies used in the study and their dilutions are indicated in the table below-

| Antibody | Source | Dilution (IB- immunoblotting, IF- immunofluorescence) |
| --- | --- | --- |
| **Primary antibodies** |  |  |
| Rabbit polyclonal α-FAM127B/RTL8A (detects all RTL8 proteins) | Proteintech #20282-1-AP | 1:500 (IB) |
| Custom made rabbit polyclonal α-FAM127B/RTL8A (detects all RTL8 proteins) | GenScript (immunogen sequence: MEGQGKVKRPKAYMC) | 1:250 (IB), 1:100 (IF) |
| Rabbit polyclonal α-FLAG | Millipore Sigma #F7425 | 1:1000 (IF) |
| Mouse monoclonal α-FLAG | Millipore Sigma #F1804 | 1:2000 (IB) |
| Rat monoclonal α-FLAG Alexa Fluor<sup>TM</sup> 488/555-conjugated | Thermo Fisher # MA1142-A488/MA1142-AF555 | 1:200 (IF) |
| Mouse monoclonal α-GAPDH | Millipore Sigma #MAB374 | 1:5000 (IB) |
| Rabbit polyclonal α-HA | Santa Cruz #sc-805 | 1:500 (IF) |
| Mouse monoclonal α-HA | BioLegend #901513 | 1:1000 (IF) |
| Rabbit polyclonal α-Histone H3 | Cell Signaling #4499S | 1:5000 (IB) |
| Mouse monoclonal α-Hsp70 | Abcam # ab5439 | 1:2000 (IB), 1:1000 (IF) |
| Mouse monoclonal α-NPM1 | Thermo Fisher #32-5200 | 1:1000 (IF) |
| Guinea pig polyclonal α-p62/SQSTM1 | Progen #GP62-C | 1:500 (IF) |
| Mouse monoclonal α-PML Alexa Fluor<sup>TM</sup> 647-conjugated (detects all PML isoforms) | Santa Cruz #sc-377390 AF647 | 1:50 (IF) |
| Rabbit polyclonal α-TDP-43 | Proteintech #10782-2-AP | 1:200 (IF) |
| Rabbit polyclonal α-UBQLN1/2 | Novus Biologicals #NBP1-56536 | 1:1000 (IB) |
| Mouse monoclonal α-UBQLN2 | Novus Biologicals #NBP2-25164 | 1:2000 (IB), 1:1000 (IF) |
| Mouse monoclonal α-A1Up (UBQLN4) | Santa Cruz #sc-136145 | 1:2000 (IB) |
| Mouse monoclonal α-Ubiquitin | Santa Cruz #sc-8017 | 1:250 (IF) |
| Rabbit polyclonal α-Ubiquitin | Cell Signaling #3933S | 1:1000 (IB) |
| **Secondary antibodies** |  |  |
| Goat α-mouse IgG(H+L) HRP-conjugated | Jackson ImmunoResearch #115-035-146 | 1:2000 (IB) |
| Goat α-rabbit IgG(H+L) HRP-conjugated | Jackson ImmunoResearch #111-035-144 | 1:2000 (IB) |
| Goat α-mouse IgG(H+L) Alexa Fluor<sup>TM</sup> 488/594/647-conjugated | Thermo Fisher #A11001/ A11004/ A21235 | 1:1000 (IF) |
| Goat α-rabbit IgG(H+L) Alexa Fluor<sup>TM</sup> 488/594/647-conjugated | Thermo Fisher #A11008/A11037/A21244 | 1:1000 (IF) |
| Goat α-guinea pig IgG(H+L) Alexa Fluor™<br>647-conjugated | Thermo Fisher #A21450 | 1:1000 (IF) |

### Plasmids

The pCMV4-FLAG-UBQLN2 plasmid (p4455 FLAG-hPLIC-2; Addgene plasmid # 8661) and pCS2-FLAG-UBQLN1 plasmid (p4458 FLAG-hPLIC-1; Addgene plasmid # 8663) were gifts from Dr. Peter Howley [65]. UBQLN4 was cloned from pDONR223-UBQLN4 (pENTR-A1UP; Addgene plasmid # 16170), which was a gift from Dr. Huda Zoghbi (Baylor College of Medicine; [66]), into the pCMV4-FLAG vector. Flag-tagged UQBLN2 deletion constructs Flag-UBQN2-ΔUBA and Flag-UBQLN2-ΔUBL were subcloned from iRFP-UBQLN2 deletion constructs described previously [22]. The UBQLN2 L619A mutation was introduced into the Flag-UBQLN2 backbone using site directed mutagenesis as described previously [22]. Mouse RTL8A cDNA was PCR amplified from first stand cDNA synthesized from RNA isolated from wildtype C57BL/6 mouse brain using primers 5’-ATGGAAGGCCAAGGCAAGGTAAAG-3’ and 5’-CTAGAAGTCCTCATCCTCCTCCCACCCGAAC-3’ and cloned into the pCMV-HA vector downstream of the HA tag using restriction sites EcoRI and KpnI. For recombinant protein studies, mRTL8A was subcloned into the pET-28 vector backbone downstream of the His-tag using restriction sites BamHI and HindIII. GST-tagged UBQLN2 full length and deletion constructs were subcloned from iRFP-UBQLN2 constructs described previously [22] into the pGEX4T vector downstream of the GST tag.

### Recombinant Protein Expression

UBQLN2 and mRTL8A constructs were transformed in Rosetta (DE3) *Escherichia coli*. All Luria-Bertani (LB)/agar plates and LB media were supplemented with 50 μg/ml kanamycin and 34 μg/ml chloramphenicol. Transformed cells were grown overnight at 37°C on LB/agar plates. On the following day, cells were transferred to 100 mL LB starter cultures and allowed to grow for 60-90 min. The starter cultures were then transferred to 2L flasks containing 1 L LB media. After the OD reached A600 ≈ 0.6-0.8, cells were induced with 0.5 mM isopropyl β-D-1-thiogalactopyranoside (IPTG) and collected after 3-5 additional hours of incubation. The bacteria were collected by centrifugation for 6 min at 10,322 x g. Bacterial pellets were stored at -80°C and purified within 1-2 days.

### Transgenic Mice

UBQLN1^-/-^ mice were generated through the University of Michigan Transgenic Animal Core. CRISPR/Cas9 was used to delete UBQLN1 exons 2 and 3 which introduces a premature stop codon after splicing from exon 1 to 4. This ablates protein expression by nonsense mediated mRNA decay. Primers for genotyping for UBQLN1: UBQLN1^-/-^ FWD – 5’-CTT GGC AAT AGG CAT TGA ATG AAA GAA GT-3’, UBQLN1^-/-^ REV – 5’-CAT CTC TTA ACC AAT GAG CTG TCT CTC C-3’ and UBQLN1 WT REV – 5’-GTA CAG TAC CAC CCA GAC TG-3’. Wild type mice for UBQLN1 amplify a band at 350 bp. Homozygous knockout mice generate a band at 550 bp and heterozygote mice amplify two bands, one at 350bp and another at 550bp. UBQLN4^-/-^ mice were a generous gift from Dr. Huda Zoghbi (Baylor College of Medicine). UBQLN4^-/-^ mice were generated from ES cell line XG905 (RRID: CVCL_PW18) (BayGenomics) created with a gene trap vector containing a splice-acceptor sequence upstream of the reporter gene, β-geo (a fusion of β-galactosidase and neomycin phosphotransferase II) splice. 5’ RACE sequencing identified the insertion site of the gene trap vector downstream of Exon 5 of the UBQLN4 gene. UBQLN4^-/-^ mice are genotyped using primers UBQLN4 β-geo FWD – 5’-ATC TTC CTG AGG CCG ATA-3’ and UBQLN4 β-geo Rev 5’-GTC AAA TTC AGA CGG CAA AC-3’ which prime within the β-geo reporter gene and amplify a band of 250 bp. β-geo positive mice are further determined to be either UBQLN4 KO heterozygotes or homozygotes by Western blot of brain lysate probed for the absence of the UBQLN4 protein. UBQLN2 WT-low and UBQLN2^-/-^ mice and their genotyping were used as previously described [22, 24].

### Cell culture and transient transfection

HEK293 (ATCC CRL-1573; Batch # 70008735) were maintained in DMEM (HyClone™ DMEM/High Glucose +L-Glutamine, +Glucose, -Sodium Pyruvate, Cytiva Life Sciences) supplemented with 10% fetal bovine serum (Atlanta Biologicals) and 1X Penicillin/Streptomycin (Invitrogen). HEK293 T-rex control and UBQLN1, UBQLN2, and UBQLN4 knock out (TKO) cells were kindly provided by Dr. Ramanujan Hegde (Medical Research Council (MRC) Laboratory of Molecular Biology; [67]) and maintained in DMEM supplemented with 10% fetal bovine serum, 10 μg/mL Blasticidin S (Fisher), and 100 μg/mL Hygromycin B (Fisher). All lines were maintained and passaged in 5% CO_2_. All described plasmids were transfected into cells using FuGENE® HD Transfection Reagent (Promega) according to the manufacturers protocol. Briefly, media was replaced with fresh complete DMEM prior to adding FuGENE/DNA mix to cells. FuGENE and DNA mixes were separately prepared in Opti-MEM® I(1X) Reduced Serum Medium (Gibco) while maintaining a FuGENE to DNA ratio of 3:1 and incubated for 5 minutes at room temperature. FuGENE/DNA master mix was prepared by adding FuGENE to DNA. This mixture was incubated for 30 minutes at room temperature prior to addition to cells.

### Immunoblotting

Protein samples in 1X SDS loading buffer/100 mM dithiothreitol (DTT) were resolved by gel electrophoresis on acrylamide NuPAGE Novex 4-12% Bis-Tris Protein Gels (Invitrogen) with NuPAGE MES SDS Running Buffer (Invitrogen). Gels were subsequently transferred onto 0.2 μM nitrocellulose membranes at 100V for 1 hour. Membranes were immediately rinsed with deionized H2O, stained with Ponceau S and imaged on a G Box Mini imager (Syngene). Membranes were rinsed with 1X Tris-Buffered Saline, 0.1% (TBST) for 10 minutes, and blocked with 5% non-fat dry milk (DotScientific) and 0.05% BSA (Fisher) in 1X TBST for 1 hour. Membranes were incubated overnight with shaking in the cold with primary antibodies diluted with 5% milk and 0.05% BSA in 1X TBST). Membranes were rinsed three times with 1x TBST for 10 minutes, incubated at room temperature with secondary antibodies for 1 hour, and then rinsed three times with 1X TBST for 10 minutes prior to developing with Western Lightning® Plus ECL (Fisher) or EcoBright Nano/Femto HRP 50 (Innovative Solutions) using the G Box Mini imager set to ECL auto exposure. GeneSys (Syngene) was used for blot quantification and raw values were normalized to their respective loading controls.

### Protein purification

#### For WT and L619A UBQLN2-ubiquitin pulldown

His-UBQLN2 pellets from WT and L619A UBQLN2 expressing *E*.*coli* were resuspended in pre-chilled lysis buffer containing 25 mM sodium phosphate, 0.5 M sodium chloride, 20 mM imidazole pH 7.4, cOmplete^™^ Mini EDTA-free Protease Inhibitor Cocktail (Sigma Aldrich; 1 tablet per 10 ml lysis buffer), 6 μL/mL of saturated phenylmethylsulfonyl fluoride (PMSF) in ethanol, 1 mg/ml lysozyme, and 10% glycerol. Bacteria were then lysed using EmulsiFlex B-15 high pressure homogenizer (Avestin). The lysate was centrifuged at 31,000 x g for 25 min. The proteins were precipitated from the supernatant by adding 0.2 g/mL ammonium sulfate. The solution was stirred at 4°C for 30 min and centrifuged at 13,000 x g for 25 min. The pellets were re-dissolved in buffer containing 25 mM sodium phosphate, 0.5 M sodium chloride, 20 mM imidazole pH 7.4, filtered through 0.22 µm Steriflip vacuum filters (Millipore) and subsequently loaded on a HisTrap HP column (GE Healthcare Life Sciences) using 25 mM sodium phosphate, 0.5 M sodium chloride, 20 mM Imidazole pH 7.4 as the binding buffer. The protein was eluted by running a 0-100% gradient of 25 mM sodium phosphate, 0.5 M sodium chloride, 0.5 M Imidazole pH 7.4. Eluates were assessed on Western blot and probed for UBQLN2 and ubiquitin.

#### For GST-UBQLN2, His-mRTL8A pulldown

His-tagged mRTL8A bacteria pellets were resuspended in pre-chilled lysis buffer containing and lysed as described above. The lysates from the EmulsiFlex B-15 homogenizer were added to Ni-NTA agarose (Qiagen) and incubated on the nutator at 4 °C for 1 h. Then the beads with the bound protein were washed twice with wash buffer (1x TBS, 10 mM imidazole, pH 8.0). Beads were then mixed with of elution buffer (1x TBS, 250mM imadizole, pH 8.0) and incubated on a nutator for 30 min at 4 °C. The slurry was spun at 700 x g for 3 min and the supernatant with the eluted protein was collected.

### Mouse brain dissection

All animal procedures were done in accordance with the Institutional Animal Care and Use Committee (IACUC) standards at the University of Michigan. Animals were deeply anesthetized with a ketamine/xylazine mixture and perfused transcardially with 0.1M phosphate buffer. Brains were dissected and lysates were prepared in RIPA buffer (cat #R0278, Sigma) supplemented with cOmplete™ Mini EDTA-free Protease Inhibitor Cocktail (Sigma Aldrich), saturated PMSF in ethanol, and Pho sSTOP™(Sigma Aldrich). Brain tissue was homogenized in a Potter homogenizer, centrifuged (13000 rpm for 30 minutes) and the supernatants were collected. For immunoblotting, protein concentration was measured using a BCA assay (cat#23227, Thermo Scientific). Samples were prepared in 1x Laemmli sample buffer before loading or storing at -20 degrees for later use. For immunoprecipitation experiments, brain lysates were used the same day as they were prepared.

### Immunoprecipitation

GST-tagged UBQLN2 bacteria pellets (Full length, ΔUBA and ΔUBL) pellets were resuspended in pre-chilled lysis buffer and lysed. Lysates were applied to GST agarose column (Glutathione agarose resin, GoldBio Cat# G-250-5) and washed 3 times in wash buffer 1x TBS + 1mM DTT). Purified His-mRTL8A was applied to column and incubated at 4°C. Column was washed 3 times in wash buffer (1x TBS + 1mM DTT) and eluted with 10mM reduced glutathione in wash buffer. Eluates were evaluated by immunoblotting and probed for UBQLN2 and RTL8. Brain lysates from transgenic UBQLN2-low mice [22] were pre-cleared with 0.1 volume of a protein A agarose beads slurry (ThermoFisher Cat# 20334) rotating at 4°C for 1 hour. The lysate was centrifuged at 4,000rpm at 4°Cfor 30 minutes and the supernatant (pre-cleared lysate) was removed. The pre-cleared lysate was incubated with anti-FLAG M2 affinity gel beads (250uL) (Sigma Cat# A2220) rotating at 4°C for 3 hours. The beads were washed 3 times with 150uL volume of PBS for 20 min each centrifuged at 4,000rpm at 4°C and the PBS was discarded. Proteins were eluted by incubating the anti-Flag beads in 2x SDS loading buffer/100mM DTT at 100 °C for 5 minutes. The supernatant was collected after centrifugation at 4,000rpm and saved for immunoblot analysis.

### Biochemical fractionation

For biochemical fractionation experiment, 2 million HEK293 cells were used per technical replicate. Cells were first washed in ice cold 1X PBS. Whole cell lysate was prepared by sonicating one-fifth volume of the total cell pellet for 10 seconds in ice-cold 1X Pierce RIPA buffer (Invitrogen) supplemented with protease inhibitors (cOmplete™ Mini EDTA-free Protease Inhibitor Cocktail Tablet, Sigma Aldrich), incubation on ice for 10 minutes followed by centrifugation at maximum speed for 10 minutes. Biochemical fractionation was done using NE-PER extraction reagents (Thermo Scientific) supplemented with Halt Protease Inhibitor Cocktail (Thermo Scientific) as per the manufacturer’s instructions with modifications as outlined below. After obtaining the nuclear soluble fraction, the remaining pellet was washed thrice in ice cold PBS. 1X RIPA buffer was added to the pellet and sonicated for 10 seconds. This was termed the total insoluble fraction. 10µg of protein from each fraction as determined by a BCA assay (Thermo Scientific) was used for immunoblotting. Blots were probed using α-RTL8 and α-UBQLN2 antibodies with GAPDH and Histone H3 serving as cytoplasmic and nuclear markers, respectively.

### Immunofluorescence

Cells were seeded onto 12mm Poly-D-lysine coated coverslips (Neuvitro Corporation) 24 hours prior to transfected with pCS2-FLAG-UBQLN1, pCMV4-FLAG-UBQLN2, pCMV4-FLAG-UBQLN4, pCMV-HA-mRTL8A and/or pCMV-HA (empty vector) using FuGENE. FuGENE/DNA mix was replaced with fresh supplemented DMEM 6 hours post transfection. 24 hours post transfection, cells were washed with 1X PBS and fixed with 4% PFA in 1X PBS. Excess PFA was neutralized using 50 mM glycine in PBS. Coverslips were then permeabilized and blocked in 10% normal goat serum (Vector Labs) in 1X PBS containing 0.1% Triton X-100 (Fisher) and 0.05% BSA (PBTGS) and incubated with primary antibodies diluted in PBTGS overnight at 4°C with shaking. For primary antibodies not conjugated to fluorophores, coverslips were washed in PBS prior to incubation with secondary antibodies (diluted in PBTGS) for 1 hour at room temperature with shaking. After washing three times in PBS, coverslips were mounted on glass slides using Prolong Glass (Invitrogen), sealed with Covergrip sealant (Biotium) and allowed to cure for 24 hours prior to imaging. Z-stack images were acquired on a Leica TCS SP5 confocal microscope using an Airyscan detector and a 100x oil objective or a Nikon A1 confocal microscope using a 63x oil objective. Final representative images were created on Adobe Photoshop.

### Quantification of colocalization, pixel intensity plot generation and measurement of nuclear-cytoplasmic ratio

To quantify colocalization, Mander’s correlation coefficients on a region of interest were obtained using the Just Another Colocalization Plugin (JACoP) [68] on ImageJ (NIH, v1.53c) with threshold value set to 20 across all channels. Statistical significance was ascertained using a Student’s t-test or one-way ANOVA as applicable. For generating pixel intensity plots, maximum intensity projections of z-stacks were generated on ImageJ. An ROI was specified using the line tool across each channel and the plot profile feature was used. The raw values obtained from each channel were normalized to the maximum value of the particular channel and plotted as a function of distance in µm. Quantification of nuclear UBQLN1, 2 and 4 was performed using Cell Profiler, an open-source cell imaging analysis software (https://cellprofiler.org/) [69]. Nuclear fluorescence intensity of UBQLN proteins was quantified by measuring the degree of colocalization of FLAG-tagged UBQLN with the DAPI signal. Cytoplasmic fluorescence intensity was measured by determining ROI based on UBQLN staining excluding the nucleus. The ratio of the nuclear to cytoplasmic fluorescence intensity was plotted across different UBQLNs in the presence and absence of overexpressed mRTL8.

### Assessment of RTL8 levels in UBQLN knockout HEK293 cells

HEK293 T-rex control (control) and UBQLN1,2,4, total knock out (TKO) cells were plated on 6-well plates. In case of rescue experiments, cells were transfected with either pCS2-FLAG-UBQLN1, pCMV4-FLAG-UBQLN2, pCMV4-FLAG-UBQLN4 or pCMV-HA (empty vector) and harvested 48 hours post transfection. Cells were washed with 1X PBS and lysed for 5 minutes in cold 1X Pierce RIPA Buffer supplemented with PMSF in ethanol and cOmplete™ Mini EDTA-free Protease Inhibitor Cocktail. Samples were sonicated and centrifuged at 21,920 x g for 30 minutes at 4°C. Soluble supernatant fractions were collected and protein concentration was determined by the Pierce BCA assay (Thermo Scientific). 20µg of protein from all samples were used for immunoblotting.

### Alignment and sequence comparison

Human and mouse RTL8 sequences were obtained from UniProt (Accession IDs – Q9BWD3 (hRTL8A), Q17RB0 (hRTL8B), A6ZKI3 (hRTL8C), Q9D1F0 (mRTL8A/B) and Q9D6I0 (mRTL8C)). Alignment was conducted with the ESPript 3.0 web tool (http://espript.ibcp.fr) [70]. Percentage identity matrix based on sequence identity was generated using Clustal Omega. (https://www.ebi.ac.uk/Tools/msa/clustalo).

### Statistical analysis

All statistical tests were carried out on GraphPad Prism 8 software (Graphpad Software Inc.). The significance threshold was set at p= 0.05, unless indicated otherwise. Details regarding specific statistical analysis are included in the figure legends.

## Declarations

### Availability of data and material

- The authors confirm that the data supporting the findings in this are available within the article, at repository links provided within the article, and within its supplementary files.
- The authors agree to share reagents, cell lines and animal models used in this study upon request.

### Code availability

Not Applicable

### Authors’ contributions

Conceptualization: HM, AP, HP, LS

Methodology: HM, AP, HT, CZ, XZ, NS, VB, LS

Investigation: HM, AP, HT, CZ, XZ, EC, RP, NS, LS

Writing: HM, HP, LS

Funding Acquisition: LS, HP

Resources: LS, YZ, HP

Supervision: HP, YZ, LS

### Ethics approval

All animal experiments were conducted under the approval of The University of Michigan Institutional Animal Care & Use Committee, protocol number PRO00010103

### Consent to participate

Not Applicable

### Consent for publication

Not Applicable

## Acknowledgments

We thank Peter Howley and Huda Zoghbi for providing constructs, Huda Zoghbi for providing the UBQLN4 -/- mouse line and Ramanujan Hegde for providing the ubiquilin triple knockout cell line. We also thank Magdalena Ivanova and Sami Barmada for their helpful suggestions and critical feedback.

---

**Supplementary Figure 1.**
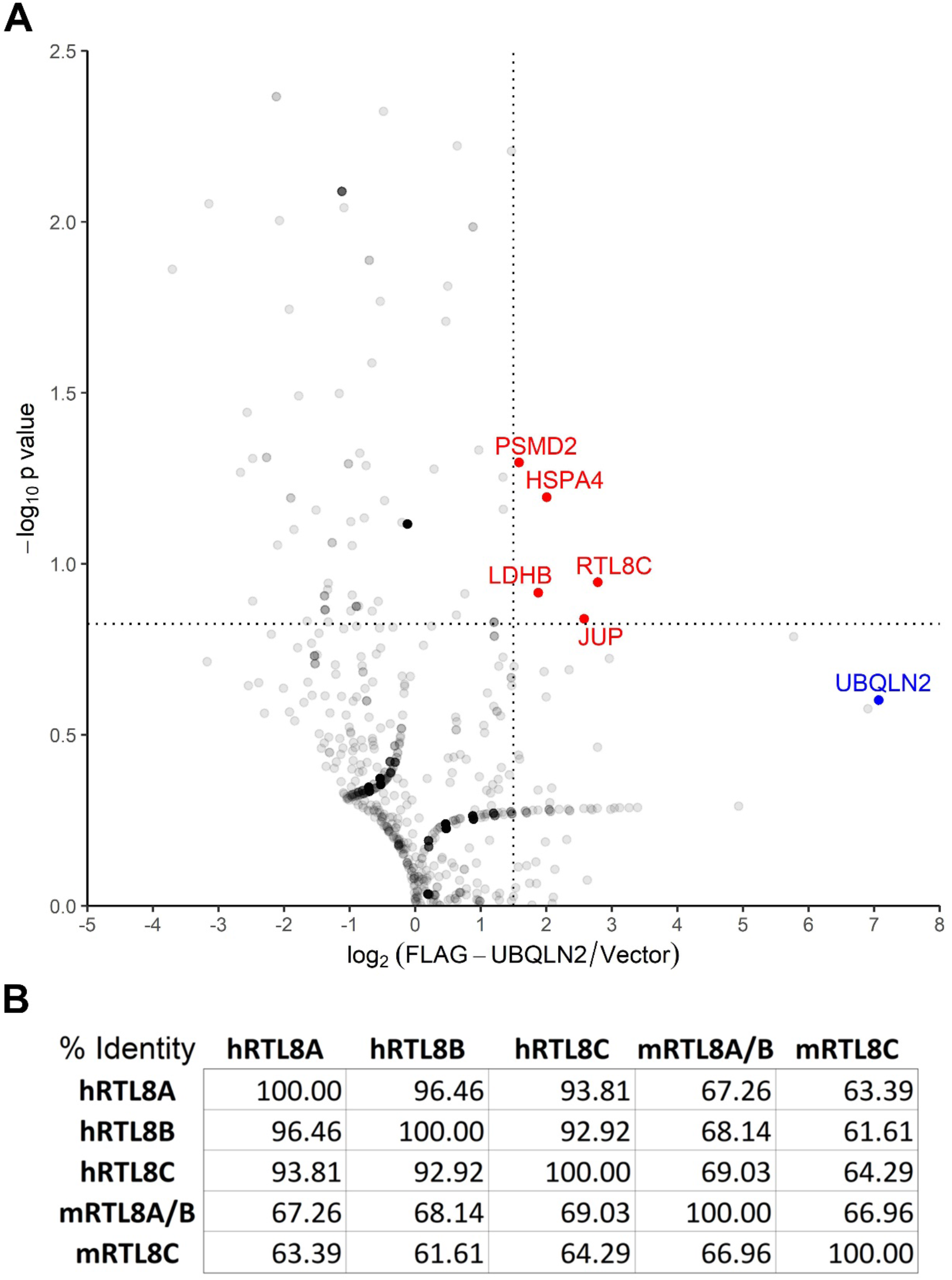
**A)** Volcano plot depicting enrichment of RTL8C pulled down from an immunoprecipitation-mass spectrometry screen in HEK293 cells identifying novel UBQLN2 interactors. A Student’s t-test was used, -log_10_ p-value versus log_2_ fold enrichment values over control plotted on the x-axis. Values with a -log_10_ p-value greater than 0.8239 (corresponding to a p-value of 0.15) and log_2_ fold enrichment greater than 1.5 are shown in red. The p-value threshold was lowered so as not to omit possible interactors since only two replicates were used for plot generation. UBQLN2 is shown in blue. **B)** Percentage Identity Matrix showing the RTL8 amino acid sequence percentage similarities for human (hRTL8) and mouse (mRTL8) proteins.

**Supplementary Figure 2.**
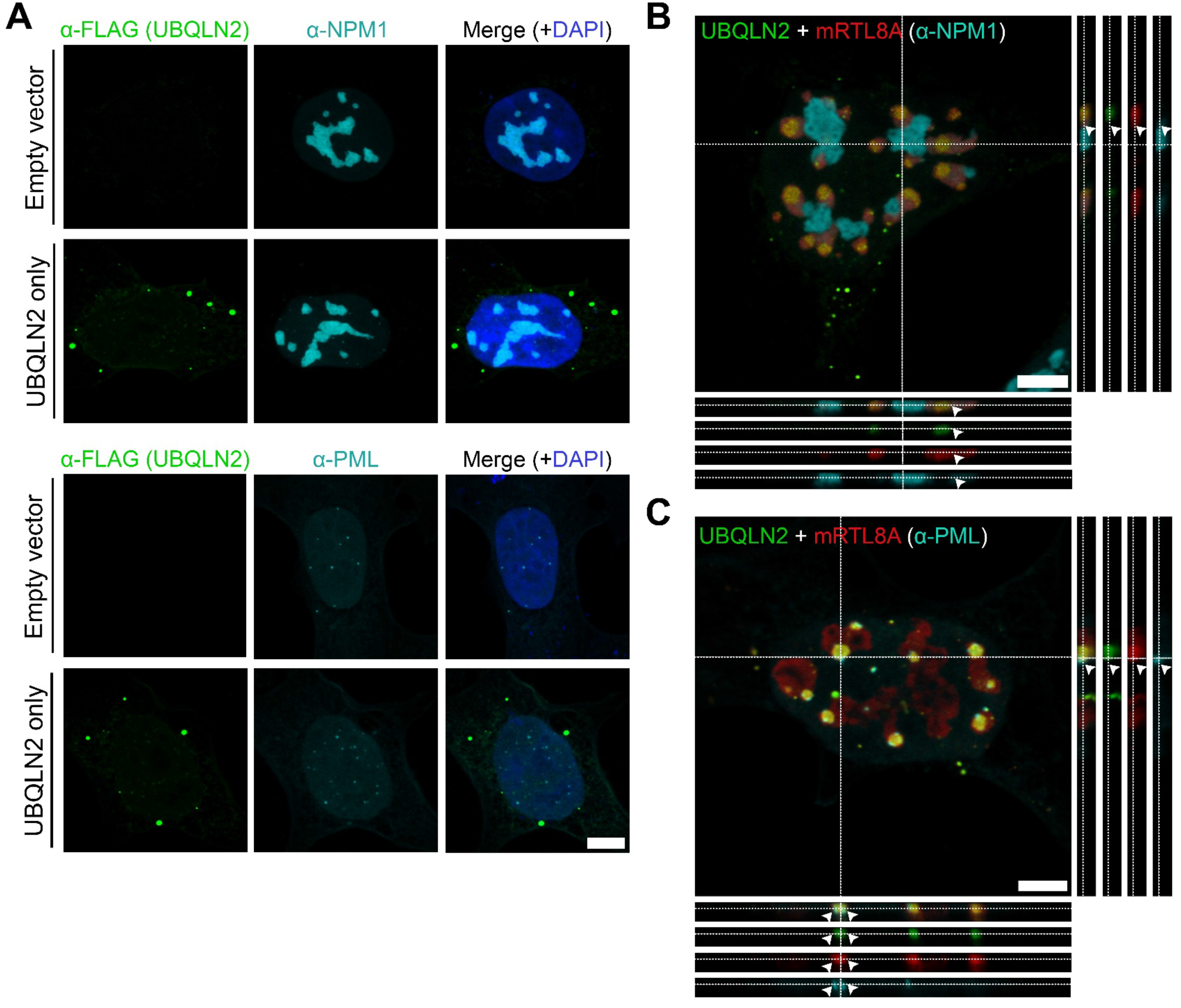
**A)** Representative images of HEK293 cells transfected with empty vector or vector encoding FLAG-UBLQN2 and stained for NPM1 or PML. (Scale bar = 5 µm). **B)**, **C)** Separate channels x-z and y-z projections for representative z stacks shown in Figure 2A and 2B. White arrows indicate UBQLN2-mRTL8A subnuclear structures that are distinct from nucleoli (NPM1) and are in close proximity (but not completely colocalizing) with PML bodies. (Scale bar = 5 µm).

**Supplementary Figure 3.**
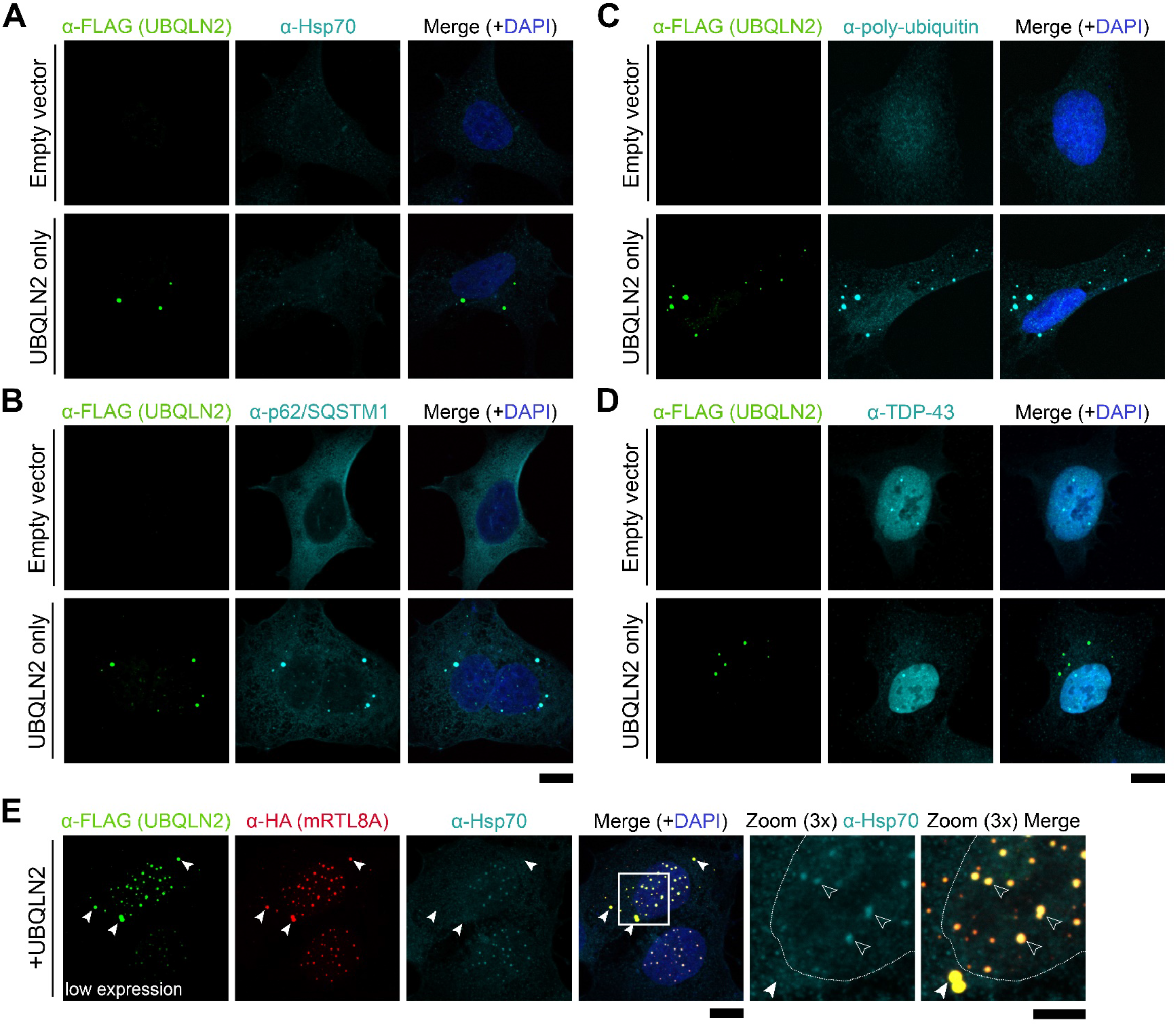
Representative images of HEK293 cells transfected with empty vector or vector encoding FLAG-UBLQN2, and stained for **A)** Hsp70, **B)** p62, **C)** poly-ubiquitin or, **D)** TDP-43. (Scale bar = 10 µm). **E)** Representative images of HEK293 cells transfected with vectors encoding FLAG-UBLQN2 and HA-mRTL8A, stained for Hsp70. In these images, nuclear mRTL8A structures (shown by unfilled arrows in the zoomed images) are not coalesced as they are in Figure 3A yet co-localize with UBQLN2 and Hsp70. Filled arrows indicate cytoplasmic UBQLN2-mRTL8A structures that do not co-localize with Hsp70. White outlines in the zoomed images represent the nucleus as determined by the extent of DAPI staining (blue in merged images). (Scale bar for merged images = 10 µm and zoomed images= 5 µm).

## References

1. Saeki, Y., et al., Ubiquitin-like proteins and Rpn10 play cooperative roles in ubiquitin-dependent proteolysis. Biochemical and Biophysical Research Communications, 2002. 293(3): p. 986–992.

2. Massey, L.K., et al., Overexpression of ubiquilin decreases ubiquitination and degradation of presenilin proteins. Journal of Alzheimer’s Disease, 2004. 6: p. 79–92.

3. Seok Ko, H., et al., Ubiquilin interacts with ubiquitylated proteins and proteasome through its ubiquitin-associated and ubiquitin-like domains. FEBS Letters, 2004. 566(1-3): p. 110–114.

4. Kaye, F.J., et al., A family of ubiquitin-like proteins binds the ATPase domain of Hsp70-like Stch. FEBS Letters, 2000. 467(2-3): p. 348–355.

5. N’Diaye, E.N., et al., PLIC proteins or ubiquilins regulate autophagy-dependent cell survival during nutrient starvation. EMBO Rep, 2009. 10(2): p. 173–9.

6. Rothenberg, C., et al., Ubiquilin functions in autophagy and is degraded by chaperone-mediated autophagy. Human Molecular Genetics, 2010. 19(16): p. 3219–3232.

7. Alexander, E.J., et al., Ubiquilin 2 modulates ALS/FTD-linked FUS-RNA complex dynamics and stress granule formation. Proc Natl Acad Sci U S A, 2018. 115(49): p. E11485–E11494.

8. Zhang, D., S. Raasi, and D. Fushman, Affinity Makes the Difference: Nonselective Interaction of the UBA Domain of Ubiquilin-1 with Monomeric Ubiquitin and Polyubiquitin Chains. Journal of Molecular Biology, 2008. 377(1): p. 162–180.

9. Dao, T.P., et al., Ubiquitin Modulates Liquid-Liquid Phase Separation of UBQLN2 via Disruption of Multivalent Interactions. Mol Cell, 2018. 69(6): p. 965–978 e6.

10. Dao, T.P. and C.A. Castaneda, Ubiquitin-Modulated Phase Separation of Shuttle Proteins: Does Condensate Formation Promote Protein Degradation? Bioessays, 2020: p. e2000036.

11. Brettschneider, J., et al., Pattern of ubiquilin pathology in ALS and FTLD indicates presence of C9ORF72 hexanucleotide expansion. Acta neuropathologica, 2012. 123(6): p. 825–839.

12. Mori, F., et al., Ubiquilin immunoreactivity in cytoplasmic and nuclear inclusions in synucleinopathies, polyglutamine diseases and intranuclear inclusion body disease. Acta Neuropathol, 2012. 124(1): p. 149–51.

13. Zeng, L., et al., Differential recruitment of UBQLN2 to nuclear inclusions in the polyglutamine diseases HD and SCA3. Neurobiology of disease, 2015. 82: p. 281–288.

14. Yang, H., et al., PolyQ-expanded huntingtin and ataxin-3 sequester ubiquitin adaptors hHR23B and UBQLN2 into aggregates via conjugated ubiquitin. The FASEB Journal, 2018. 32(6): p. 2923–2933.

15. Gerson, J.E., et al., Ubiquilin-2 differentially regulates polyglutamine disease proteins. Human Molecular Genetics, 2020.

16. Deng, H.X., et al., Mutations in UBQLN2 cause dominant X-linked juvenile and adult-onset ALS and ALS/dementia. Nature, 2011. 477(7363): p. 211–5.

17. Le, N.T., et al., Motor neuron disease, TDP-43 pathology, and memory deficits in mice expressing ALS-FTD-linked UBQLN2 mutations. Proc Natl Acad Sci U S A, 2016. 113(47): p. E7580–e7589.

18. Picher-Martel, V., et al., Neuronal Expression of UBQLN2(P497H) Exacerbates TDP-43 Pathology in TDP-43(G348C) Mice through Interaction with Ubiquitin. Mol Neurobiol, 2019. 56(7): p. 4680–4696.

19. Chang, L. and M.J. Monteiro, Defective Proteasome Delivery of Polyubiquitinated Proteins by Ubiquilin-2 Proteins Containing ALS Mutations. PLoS One, 2015. 10(6): p. e0130162.

20. Hjerpe, R., et al., UBQLN2 Mediates Autophagy-Independent Protein Aggregate Clearance by the Proteasome. Cell, 2016. 166(4): p. 935–949.

21. Osaka, M., D. Ito, and N. Suzuki, Disturbance of proteasomal and autophagic protein degradation pathways by amyotrophic lateral sclerosis-linked mutations in ubiquilin 2. Biochem Biophys Res Commun, 2016. 472(2): p. 324–31.

22. Sharkey, L.M., et al., Mutant UBQLN2 promotes toxicity by modulating intrinsic self-assembly. Proceedings of the National Academy of Sciences, 2018. 115(44): p. E10495–E10504.

23. Halloran, M., et al., Amyotrophic lateral sclerosis-linked UBQLN2 mutants inhibit endoplasmic reticulum to Golgi transport, leading to Golgi fragmentation and ER stress. Cell Mol Life Sci, 2019.

24. Sharkey, L.M., et al., Modeling UBQLN2-mediated neurodegenerative disease in mice: Shared and divergent properties of wild type and mutant UBQLN2 in phase separation, subcellular localization, altered proteostasis pathways, and selective cytotoxicity. Neurobiology of Disease, 2020: p. 105016.

25. Wu, J.J., et al., ALS/FTD mutations in UBQLN2 impede autophagy by reducing autophagosome acidification through loss of function. Proceedings of the National Academy of Sciences, 2020: p. 201917371.

26. Rutherford, N.J., et al., Unbiased screen reveals ubiquilin-1 and -2 highly associated with huntingtin inclusions. Brain research, 2013. 1524: p. 62–73.

27. Latonen, L., et al., Proteasome inhibitors induce nucleolar aggregation of proteasome target proteins and polyadenylated RNA by altering ubiquitin availability. Oncogene, 2011. 30(7): p. 790–805.

28. Frottin, F., et al., The nucleolus functions as a phase-separated protein quality control compartment. Science, 2019. 365(6451): p. 342–347.

29. Mediani, L., et al., Defective ribosomal products challenge nuclear function by impairing nuclear condensate dynamics and immobilizing ubiquitin. The EMBO journal, 2019. 38(15): p. e101341–e101341.

30. Gallardo, P., et al., Acute Heat Stress Leads to Reversible Aggregation of Nuclear Proteins into Nucleolar Rings in Fission Yeast. Cell Reports, 2020. 33(6).

31. Frattini, A., et al., A Low-Copy Repeat in Xq26 Represents a Novel Putatively Prenylated Protein Gene (CXX1) and Its Pseudogenes (DXS9914, DXS9915, and DXS9916). Genomics, 1997. 46(1): p. 167–169.

32. Brandt, J., A.M. Veith, and J.N. Volff, A family of neofunctionalized Ty3/gypsy retrotransposon genes in mammalian genomes. Cytogenetic and Genome Research, 2005. 110(1-4): p. 307–317.

33. Boot, A., et al., Methylation associated transcriptional repression of ELOVL5 in novel colorectal cancer cell lines. PLoS One, 2017. 12(9): p. e0184900.

34. Whiteley, A.M., et al., Global proteomics of Ubqln2-based murine models of ALS. J Biol Chem, 2020.

35. Zheng, W., et al., Deep-learning contact-map guided protein structure prediction in CASP13. Proteins-Structure Function and Bioinformatics, 2019. 87(12): p. 1149–1164.

36. Zhang, C., et al., Blinded Testing of Function Annotation for uPE1 Proteins by I-TASSER/COFACTOR Pipeline Using the 2018–2019 Additions to neXtProt and the CAFA3 Challenge. Journal of proteome research, 2019. 18(12): p. 4154–4166.

37. Xu, J. and Y. Zhang, How significant is a protein structure similarity with TM-score= 0.5? Bioinformatics, 2010. 26(7): p. 889–895.

38. Welch, W.J. and J.R. Feramisco, Nuclear and nucleolar localization of the 72,000-dalton heat shock protein in heat-shocked mammalian cells. J Biol Chem, 1984. 259(7): p. 4501–13.

39. Lewis, M.J. and H.R. Pelham, Involvement of ATP in the nuclear and nucleolar functions of the 70 kd heat shock protein. Embo j, 1985. 4(12): p. 3137–43.

40. Pankiv, S., et al., Nucleocytoplasmic shuttling of p62/SQSTM1 and its role in recruitment of nuclear polyubiquitinated proteins to promyelocytic leukemia bodies. The Journal of biological chemistry, 2010. 285(8): p. 5941–5953.

41. Pankiv, S., et al., Nucleocytoplasmic shuttling of p62/SQSTM1 and its role in recruitment of nuclear polyubiquitinated proteins to promyelocytic leukemia bodies. J Biol Chem, 2010. 285(8): p. 5941–53.

42. Pikkarainen, M., et al., Distribution and pattern of pathology in subjects with familial or sporadic late-onset cerebellar ataxia as assessed by p62/sequestosome immunohistochemistry. Cerebellum, 2011. 10(4): p. 720–31.

43. Baloh, R.H., TDP-43: the relationship between protein aggregation and neurodegeneration in amyotrophic lateral sclerosis and frontotemporal lobar degeneration. The FEBS journal, 2011. 278(19): p. 3539–3549.

44. Udan-Johns, M., et al., Prion-like nuclear aggregation of TDP-43 during heat shock is regulated by HSP40/70 chaperones. Human molecular genetics, 2014. 23(1): p. 157–170.

45. Kang, Y., et al., Ubiquitin Receptor Proteins hHR23a and hPLIC2 Interact. Journal of Molecular Biology, 2007. 365(4): p. 1093–1101.

46. Marín, I., The ubiquilin gene family: evolutionary patterns and functional insights. BMC Evolutionary Biology, 2014. 14(1): p. 63.

47. Zheng, T., Y. Yang, and C.A. Castaneda, Structure, dynamics and functions of UBQLNs: at the crossroads of protein quality control machinery. Biochem J, 2020. 477(18): p. 3471–3497.

48. Samant, R.S., et al., Distinct proteostasis circuits cooperate in nuclear and cytoplasmic protein quality control. Nature, 2018. 563(7731): p. 407–411.

49. Jachimowicz, R.D., et al., UBQLN4 Represses Homologous Recombination and Is Overexpressed in Aggressive Tumors. Cell, 2019. 176(3): p. 505-519.e22.

50. Hernandez-Verdun, D., The nucleolus: a model for the organization of nuclear functions. Histochemistry and Cell Biology, 2006. 126(2): p. 135.

51. Boisvert, F.-M., et al., The multifunctional nucleolus. Nature Reviews Molecular Cell Biology, 2007. 8(7): p. 574–585.

52. Gerson, J.E., et al., Shared and divergent phase separation and aggregation properties of brain-expressed ubiquilins. Scientific Reports, 2021. 11(1): p. 287.

53. Moran, D.M., H. Shen, and C.G. Maki, Puromycin-based vectors promote a ROS-dependent recruitment of PML to nuclear inclusions enriched with HSP70 and Proteasomes. BMC Cell Biol, 2009. 10: p. 32.

54. Guo, L., et al., A cellular system that degrades misfolded proteins and protects against neurodegeneration. Mol Cell, 2014. 55(1): p. 15–30.

55. Lamoliatte, F., et al., Uncovering the SUMOylation and ubiquitylation crosstalk in human cells using sequential peptide immunopurification. Nat Commun, 2017. 8: p. 14109.

56. Rott, R., et al., SUMOylation and ubiquitination reciprocally regulate α-synuclein degradation and pathological aggregation. Proc Natl Acad Sci U S A, 2017. 114(50): p. 13176–13181.

57. Jin, J., Interplay between ubiquitylation and SUMOylation: Empowered by phase separation. J Biol Chem, 2019. 294(42): p. 15235–15236.

58. Pankiv, S., et al., Nucleocytoplasmic shuttling of p62/SQSTM1 and its role in recruitment of nuclear polyubiquitinated proteins to promyelocytic leukemia bodies. J Biol Chem, 2010. 285(8): p. 5941–53.

59. Beauclair, G., et al., JASSA: a comprehensive tool for prediction of SUMOylation sites and SIMs. Bioinformatics, 2015. 31(21): p. 3483–3491.

60. Flores, B.N., et al., An Intramolecular Salt Bridge Linking TDP43 RNA Binding, Protein Stability, and TDP43-Dependent Neurodegeneration. Cell Rep, 2019. 27(4): p. 1133-1150.e8.

61. Seyfried, N.T., et al., Multiplex SILAC analysis of a cellular TDP-43 proteinopathy model reveals protein inclusions associated with SUMOylation and diverse polyubiquitin chains. Mol Cell Proteomics, 2010. 9(4): p. 705–18.

62. Dangoumau, A., et al., Protein SUMOylation, an emerging pathway in amyotrophic lateral sclerosis. Int J Neurosci, 2013. 123(6): p. 366–74.

63. Zybailov, B., et al., Statistical Analysis of Membrane Proteome Expression Changes in Saccharomyces cerevisiae. Journal of Proteome Research, 2006. 5(9): p. 2339–2347.

64. Wickham, H., ggplot2: Elegant Graphics for Data Analysis. Ggplot2: Elegant Graphics for Data Analysis, 2009: p. 1–212.

65. Kleijnen, M.F., et al., The hPLIC Proteins May Provide a Link between the Ubiquitination Machinery and the Proteasome. Molecular Cell, 2000. 6(2): p. 409–419.

66. Lim, J., et al., A Protein–Protein Interaction Network for Human Inherited Ataxias and Disorders of Purkinje Cell Degeneration. Cell, 2006. 125(4): p. 801–814.

67. Itakura, E., et al., Ubiquilins Chaperone and Triage Mitochondrial Membrane Proteins for Degradation. Mol Cell, 2016. 63(1): p. 21–33.

68. Bolte, S. and F.P. Cordelières, A guided tour into subcellular colocalization analysis in light microscopy. J Microsc, 2006. 224(Pt 3): p. 213–32.

69. Carpenter, A.E., et al., CellProfiler: image analysis software for identifying and quantifying cell phenotypes. Genome Biology, 2006. 7(10): p. R100.

70. Robert, X. and P. Gouet, Deciphering key features in protein structures with the new ENDscript server. Nucleic Acids Research, 2014. 42(W1): p. W320–W324.

